# A computational dynamic model of combination treatment for type II inhibitors with asciminib

**DOI:** 10.1101/2025.02.21.639442

**Authors:** J. Roadnight Sheehan, Astrid S. de Wijn, Ran Friedman

**Affiliations:** Department of Mechanical and Industrial Engineering, Norwegian University of Science and Technology, Trondheim, Norway; Department of Chemistry and Biomedical Sciences, Linnaeus University, Kalmar, Sweden

## Abstract

Despite continuous strides forward in drug development, resistance to treatment looms large in the battle against cancer as well as communicable diseases. Chronic myeloid leukaemia (CML) is treated with targeted therapy and treatment is personalised when resistance arises. It has been extensively studied and is used as a model for targeted therapy. In this work, we examine combination treatments of type II Abl1 inhibitors and asciminib (an allosteric regulator) through a computational model at patient relevant concentrations. Due to the separate binding sites of type II inhibitors and asciminib, we propose their combination treatment as potentially robust to resistance. We find that the simultaneous cobinding of type II inhibitors and asciminib is high in synergetic combinations. As an aid to designing and comparing combination treatments, we put forward an equation that expands on the effective ratio of IC_50_ (ERIC). Unlike usual comparisons of IC_50_ values, ERIC takes takes patient plasma concentrations into account. The product of two ERIC values (ERIC_combo_) creates comparable approximations of the effectiveness of combination treatments with low levels of synergy or antagonism at different concentrations. Its simple formulation is done without experiments and requires less computation and input data than the current standard of ZIP values. As such, the new scheme is a useful complement to experiments that deal with synergy in drug use.

## 1 Introduction

Drug design and nature are in an “arms race”. In cancers and communicable diseases, resistance to treatment arises and one way to combat this is further steps in drug development. A blood cancer that has been extensively studied as a model for this showdown of medicine and nature is chronic myeloid leukaemia (CML) [1, 2]. Targeted therapy is used to treat CML and when resistance to this treatment develops, treatment is personalised. The level of personalisation can be high for CML patients that develop resistance to treatment as the patient’s tolerance to treatment and the type of resistance must be taken into account. This personalisation is not a straightforward process as current common methods of comparing treatment are flawed. Combination therapy is currently underutilised in this personalisation.

The frontline therapies for CML are Abl1 inhibitors. The Abl1 kinase is an important part of signalling in cells’ growth and multiplication pathways. In CML, the Abl1 kinase becomes unregulated and normal cell differentiation and death is blocked [3]. In 95% of CML cases, the unregulated Abl1 arises from a chromosomal translocation - the Philadelphia chromosome [4]. The detection of the Philadelphia chromosome is used as the basis for diagnosis. The chromosomal translocation that forms the Philadelphia chromosome is a fusion between the breakpoint cluster region (BCR) of chromosome 22 with the Ableson luekaemia (Abl) gene on chromosome 9. This fusion BCR-Abl region encodes an Abl1 kinase that lacks a key regulatory part of its structure. Not only is the Philadelphia chromosome the primary cause of CML, it is also a contributing factor in other cancers [5].

Successful treatment for CML patients is normally classed by long-term deep molecular response (DMR) and treatment free remission (TFR). The molecular response of the treatment is monitored by assessing the proportion of cell populations with the Philadelphia chromosome [6]. If there is statistically significant decrease in the molecular response, the cause of this resistance to treatment must be identified and treatment changed. Around 1 in 4 patients with CML, who are initially treated with imatinib, will develop resistance to treatment within two years [7] and in about 1 in 5 patients undertaking secondary treatment (i.e. with prior treatment of one Abl1 inhibitor) resistance will develop within one year [8]. In most of the cases, resistance develops due to mutations in the Abl1 kinase, which alter the binding affinity with the inhibitor. The mutation is identified through sequencing of the patient’s tumour.

The selection process of which Abl1 inhibitor would be best suited to treat the detected mutant Abl1 is guided by IC50 values. Unfortunately, this widely-used measure of drug efficacy is not always reliable: the variability of in vitro measurements of IC50 from different assays suggests inaccuracies, which can lead to noise in data collected from a variety of sources [9, 10, 11, 12]. Furthermore, the values examined are normally fold-IC50 values for mutants (Mut) that are relative to the wild-type (WT) Abl1 enzyme - i.e. 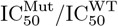. This comparison gives an answer to which mutant each drug is most potent against, but not which treatment is most potent for each mutant. In our previous work [13], we proposed an alternative parameter, Effective Ratio of IC50 (ERIC), where a measure of inhibitor concentration is included. This accounts for the differences in relative size of normal patient doses for the inhibitors compared with their IC50 values. For example, a usual dose of 400 mg of imatinib can give a concentration of roughly 2.6-fold the IC50 for the WT, whereas the usual 45 mg dose of ponatinib, can yield a concentration around 26-fold the IC50 for the WT. Without considering the context of the concentration of the inhibitor, the comparison of IC50 values could be misleading. In this work, we develop and test a model for the synergetic behaviour of combination treatments at patient-relevant concentrations of Abl1 inhibitors and we aim to demonstrate how the ERIC values can be applied in combination therapy.

The range of CML treatments to choose from is widening as new Abl1 inhibitors are being developed. There are many types of Abl1 kinase inhibitors. They are categorised by their binding site and the configuration of the kinase when binding. This work looks at three type II inhibitors (imatinib, ponatinib and nilotinib), that bind to the inactive Abl1 at the ATP site, and one type IV inhibitor (asciminib) that binds to an allosteric site far from the ATP site - i.e. the myristoyl pocket [14]. The type II inhibitors are used widely as first-line treatment of CML, but as previously stated, resistance can arise during treatment. Asciminib has proven a good tool in the campaign against type II resistant associated mutants (like those outlined in Table 1) and has a lower toxicity for patients [15]. However, asciminib has its own array of resistant Abl1 mutants. Due to the difference in binding site, there is no overlap in the resistant mutants for asciminib and type II inhibitors, meaning combination treatment could prove to be a method that is less vulnerable to the development of resistance for long term DMR in patients. This work models combination therapy as a way to prevent resistance and potentially provide a suitable, simple equation to calculate treatments at lower doses by using a type-II inhibitor in combination with asciminib.

**Table 1:** The table (data refined from [16]) shows the mutants associated with resistance for three type II inhibitors: imatinib, ponatinib, and nilotinib. These mutants were considered in this work as they are common and have been extensively studied. *Although these mutations do have an increased IC_50_ for ponatinib, they do not lead to resistance alone, but do so as compound mutations.

| Drug | imatinib | ponatinib | nilotinib |
| --- | --- | --- | --- |
| **Mutants considered in this work** | G250E<br>E255K<br>E255V<br>T315I | E255K*<br>E255V*<br>T315I* | T315I |

Interactions between different medications play an important role in prescribing treatments to patients - some interactions can be harmful and others are beneficial. In combination treatments for CML, combining two non-competitive Abl1 inhibitors could enhance treatment courses. In pharmacology, synergy describes the ability of two drugs to perform better in combination than they are expected to based on their individual abilities. The opposite effect is antagonism. If a treatment combination is synergetic, it may be possible to achieve DMR in patients with lower doses of the medication, which would benefit patients that experience more adverse side effects from treatment. This enhanced molecular response could also be applied to when CML is in acute- or blast-phase as the need for molecular response is much more time critical [17].

Computational and in vivo models of combination treatment have indicated varying levels of synergy for type II inhibitors in combination with asciminib [18, 19, 20, 21, 22, 23]. This has been mirrored by patient responses in clinical trials [24, 25]. Although this data is available, there is a lack of unity in how synergy is reported and sometimes a lack of precision in how this data is represented. This makes it challenging to plan combination treatments tailored to specific patients.

Predicting patient responses in the clinic from models’ and assays’ measurements of synergy for combination therapies is not straight-forward. While some models of treatment account for realistic in-patient concentrations of Abl1 inhibitors, cellular assays are often treated with much lower concentrations than patients. The relative size of the effects of synergy and antagonism at low doses can be very different than those at patient-relevant concentrations [18, 19, 26]. Presenting synergy data is difficult (due to its multidimensional nature) which creates many opportunities for misinterpretation. In addition to this, there is a lack of consistency in this field of how synergy is measured, despite attempts to unify this [26]. Similar to comparing IC_50_ values from different sources, comparing different levels of synergy via different measurements of synergy is difficult.

## 2 Results

The model was driven by different doses of type II inhibitors and asciminib that fluctuated to mimic the pharmacodynamics expected in patients. Each type II inhibitor (400 mg imatinib once-daily, 45 mg ponatinib once-daily and 400 mg nilotinib twice-daily) and two regimes of asciminib (80 mg once-daily and 40 mg twice-daily) were initially simulated as solo treatments at full normal doses as seen quantified in section 4.1.4. The fluctuating concentrations for 100% doses can be seen in Figure 1. As the averages of the inhibitor concentration in the steady state are similar for both regimes of asciminib, we present only once-daily doses of asciminib in the main text; further results for the twice-daily regime are offered in the supporting information.

**Figure 1.**
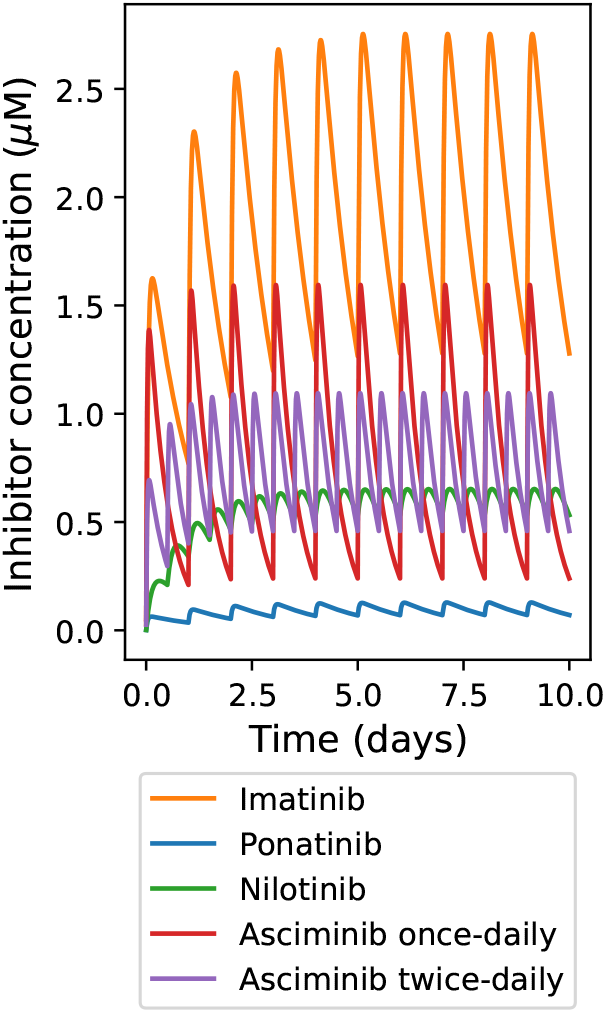
Simulated inhibitor plasma concentrations at 100% doses, calculated as outlined in section 4.1.4.

To examine the response of the system to different levels of combination therapy, other doses were selected based on the accessibility of pill-cutters - 25%, 50% and 75%. Figure 2 shows an example of the response of each state of the Abl1 enzyme to the fluctuating inhibitor concentrations. This example shows 50% doses of imatinib and once-daily asciminib for the T315I mutant enzyme. The “zoomed-in” section of this figure indicates that the individual binding of imatinib and asciminib to the active form of the enzyme play more minor parts of the inhibition process. Further examples of the varying enzyme states can be found in the supporting information (Figure S1). Figure 3 shows that cobinding of the two inhibitors for the mutant G250E enzyme is at much lower levels than with the other mutations.

**Figure 2.**
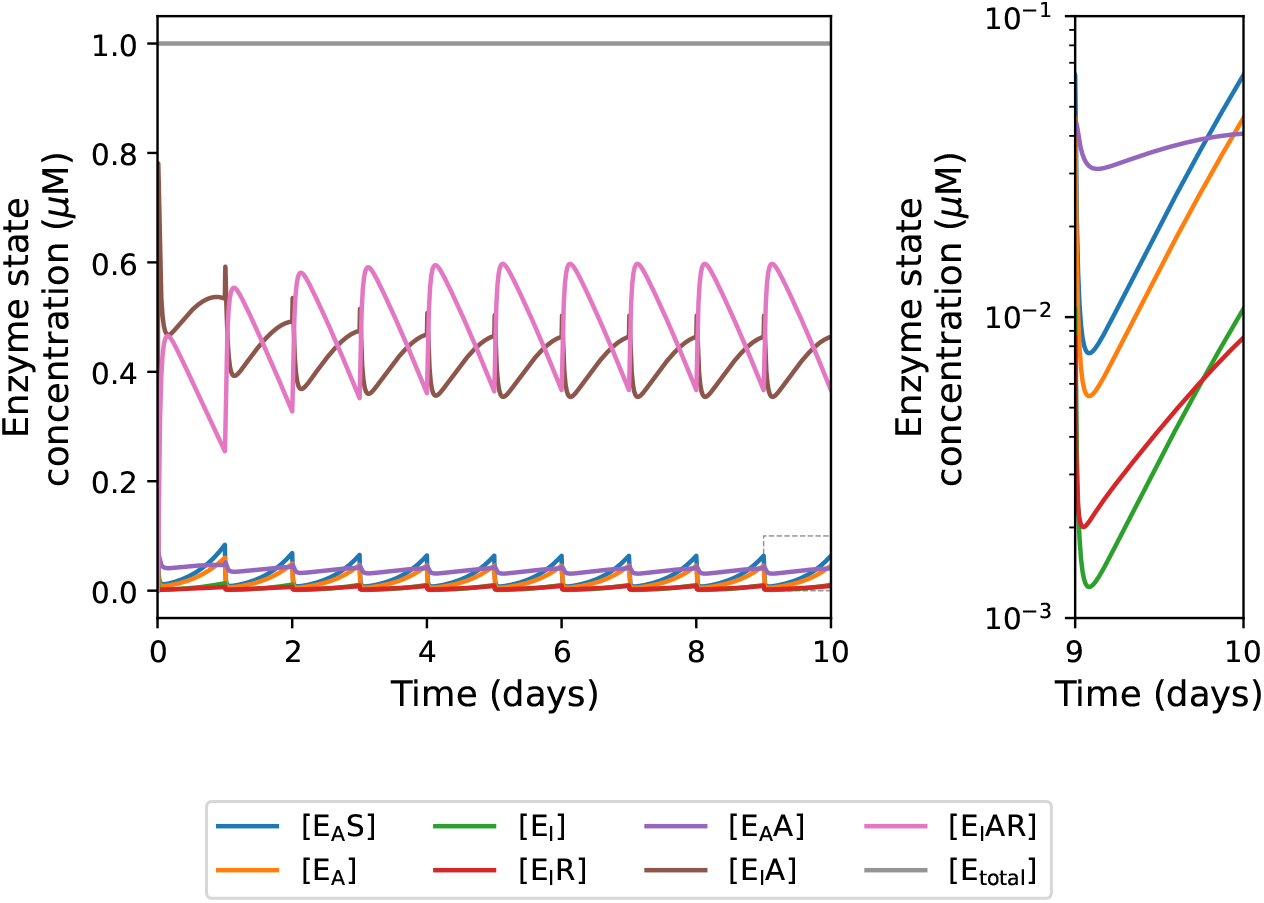
Example of an output of a 50% dose of imatinib combined with a 50% dose of the once-daily regime of asciminib for the mutant T315I enzyme. A close-up of the near-zero section on the tenth day is given to emphasise how little the single binding of imatinib (red) and asciminib to the active state (purple) contribute to this system.

**Figure 3.**
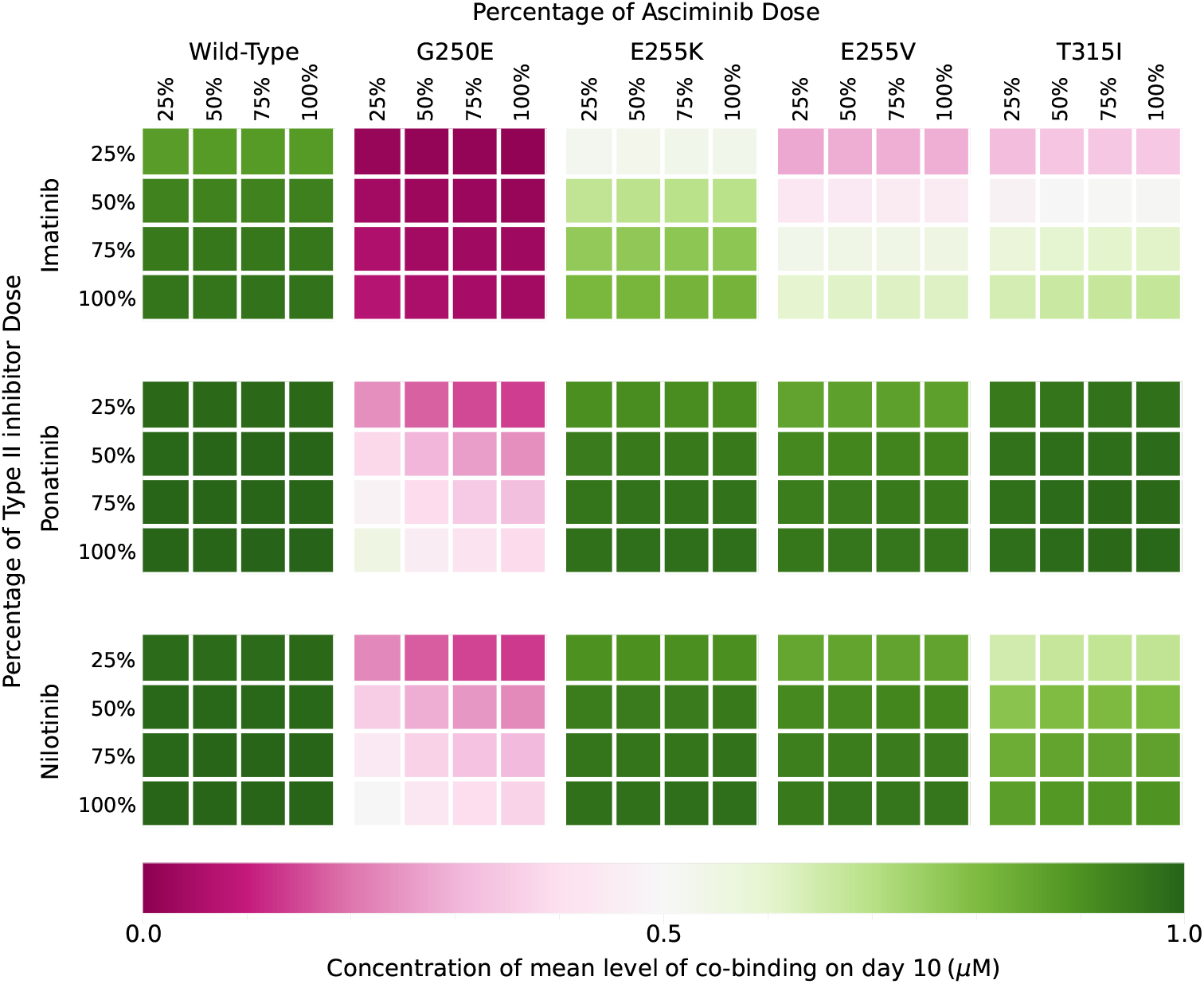
Heat map of the mean concentration of enzymes that are bound to both the asciminib and a type II inhibitor on day 10 for different combinations of doses of type II inhibitor and the once-daily regime of asciminib. Generally, we see that the co-binding is lower in systems with the G250E mutant.

As established in the previous work [13] that focused on monotherapy, the size of the product formation rate does not reveal much information to indicate resistance, but its relative decrease does. We measure this relative decrease as Inhibitory Reduction Prowess (IRP), in this work we define IRP as

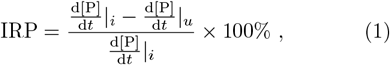

where 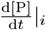 is the initial product formation rate prior to treatment and 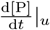 is the product formation rate that will ultimately be compared - i.e. half-height when the inhibitor concentration is in a steady-state for this work. To illustrate this concept we provide an example of product formation rates (Figure 4) of different mutants under the same treatment of 100% dose of nilotinib as monotherapy. Here it is expected that mutant T315I is associated with resistance to nilotinib; however, it does not have the highest product formation rate, showing that the absolute size of the product rate does not give indication of resistance. Figure 5 gives details of the IRP outputs for all 100% single drug treatments for the model. These give a much better indication of associated resistance than a measurement of the product rate.

**Figure 4.**
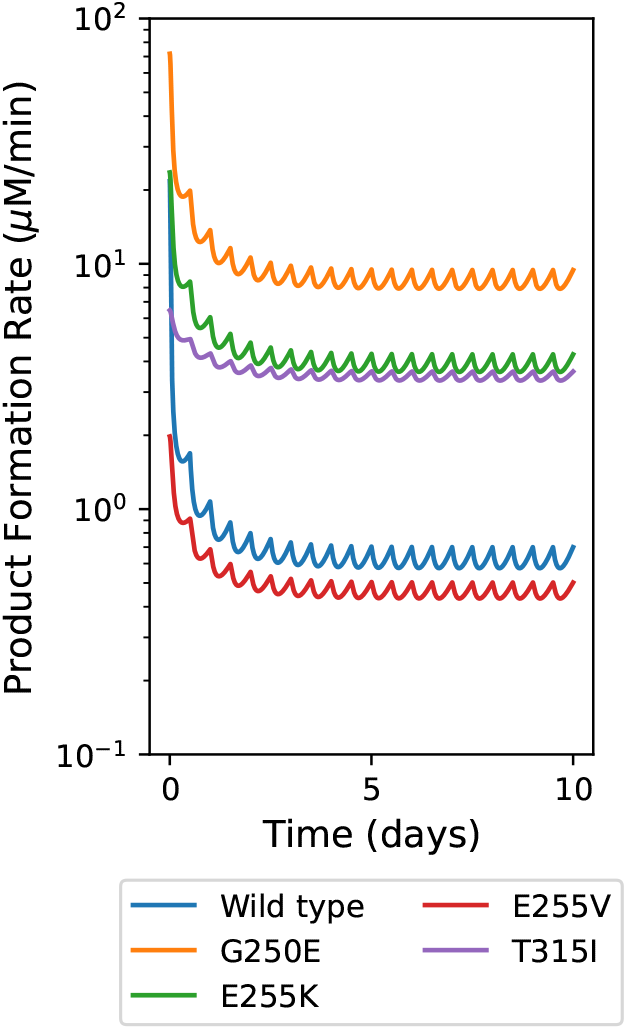
An example of the product formation rates produced in the system with 100% dose of nilotinib as monotherapy. Mutant T315I is associated with resistance to nilotinib, yet it does not have the highest product formation rate, showing that the absolute size of the product rate does not give indication of resistance.

**Figure 5.**
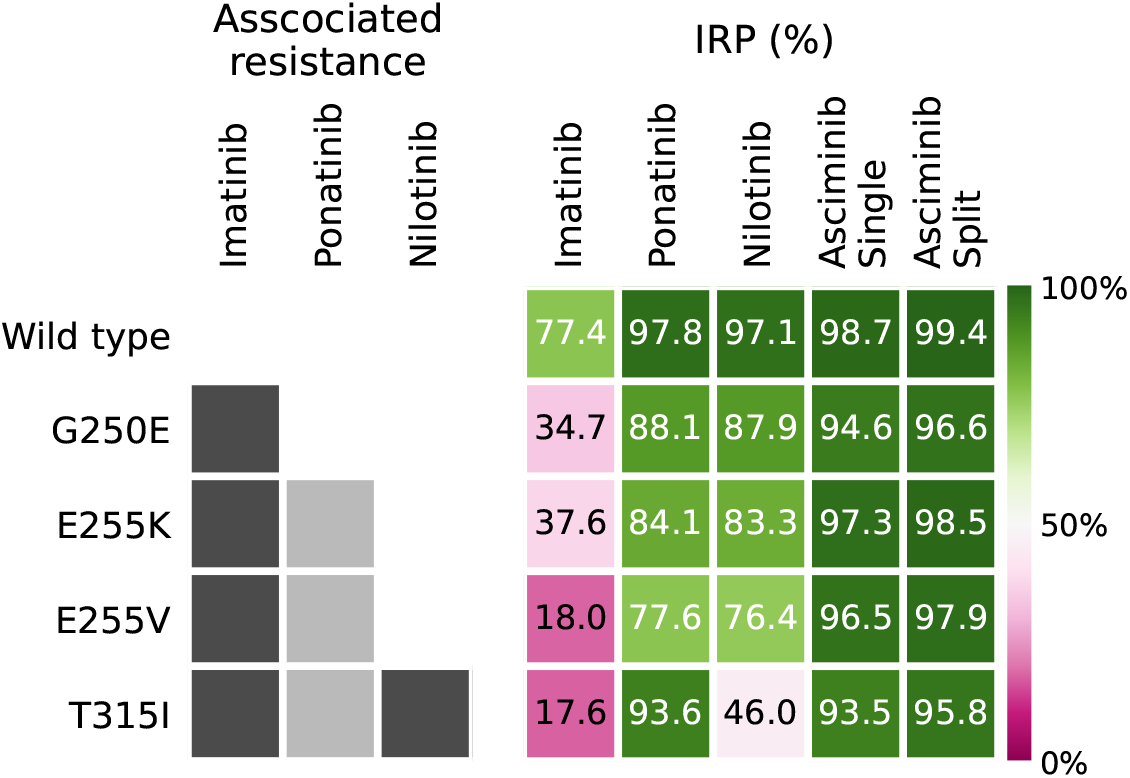
Heat map of associated resistance to type II inhibitors for mutants and the IRP values of 100% dose single drug therapy. Dark grey indicates that resistance is associated with the mutant and light grey indicates resistance associated with the mutation in combination mutations. The IRP values are for the midpoint of the range of product formation rate on day 10. The IRP values reflect the associated resistance.

To give a holistic view of how percentage of dose taken affects the IRP in combination therapy for each mutation, Figure 6 shows a heat map for each combination under the once-daily asciminib regime. Marked are indicators of IRP values of *>* 80% and *>* 95% with triangles and circles, respectively. We also present results for three asciminib resistant mutants in the supporting information (Figures S18 to S21). As the source of IC_50_ values for these mutants showed wildly different values for the WT for the type II inhibitors and asciminib (Tables S2 and S3), they are not compared directly with those results in the main text. This also reiterates the flaws in using IC_50_ values from different sources.

**Figure 6.**
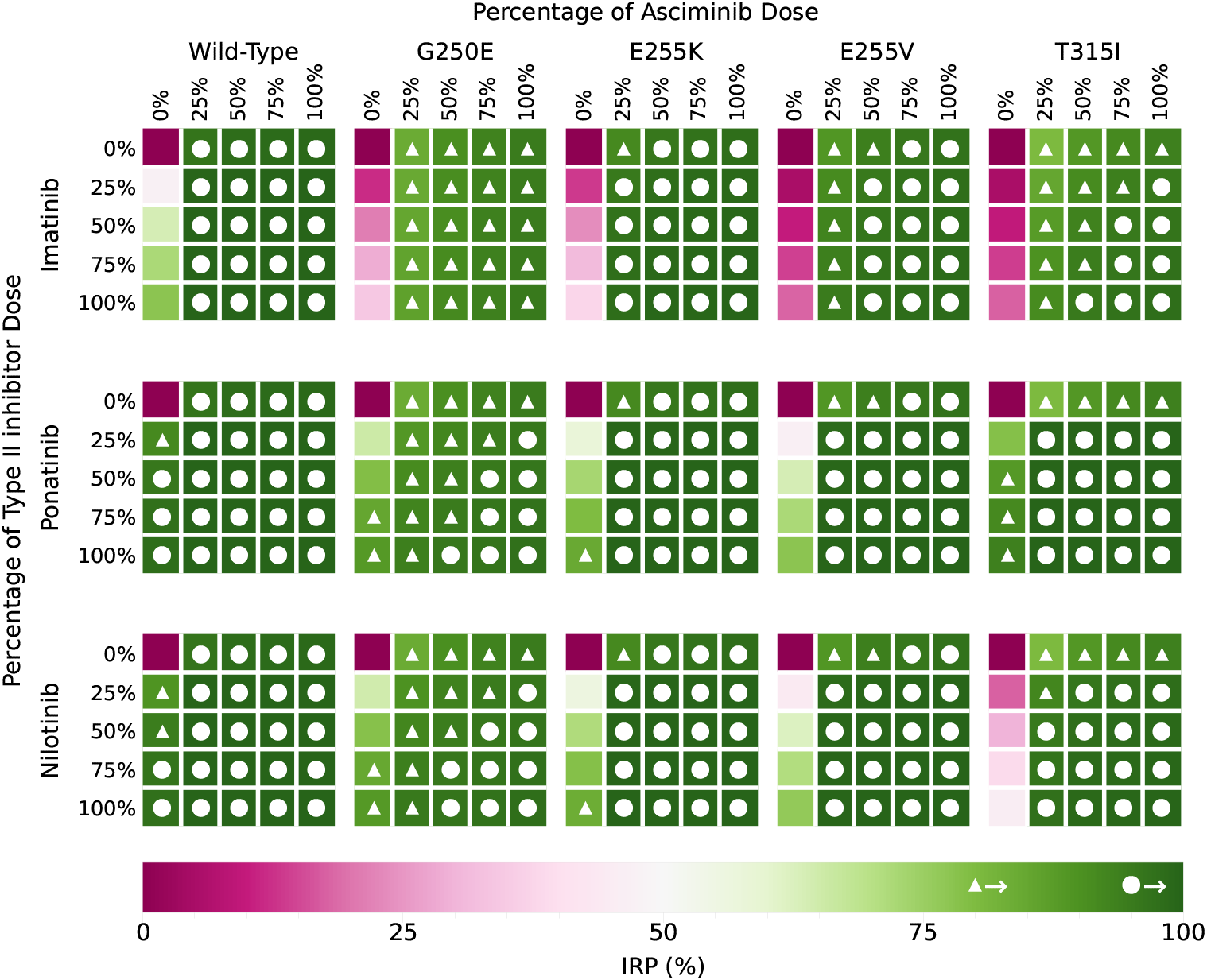
Heat map of IRP values for the midpoint of the range of product formation rate on day 10 for different combinations of doses of type II inhibitor and once-daily doses of asciminib. The white triangles and circles indicate IRPs that are over 80% and 95%, respectively. An equivalent figure for twice-daily doses of asciminib can be found in the supporting information (Figure S2), along with separate heat maps for each type II inhibitor and asciminib regime combination with individual IRP values labelled (Figures S3 to S5). This shows how different combination therapies can create similar outcomes in IRP.

As mentioned in the introduction (section 1), we aim to apply ERIC values in place of fold-IC_50_ values as means of guiding treatment selection. For solo-therapies, we define ERIC as:

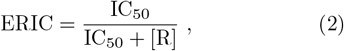

where [R] is the inhibitor concentration. To apply ERIC to combinations of inhibitor and asciminib doses for each mutant, we calculate a combined ERIC value (ERIC_combo_) as the product of each drug’s ERIC value:

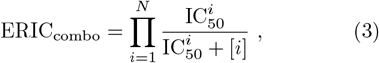

where 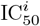 is the IC_50_ value of drug *i*, [*i*] is the concentration of drug *i*, and *N* is the total number of drugs. ERIC_combo_ is an estimate and does not account for certain aspects of the system, these will be discussed in section 3. The ERIC_combo_ values are represented in Figure 7.

**Figure 7.**
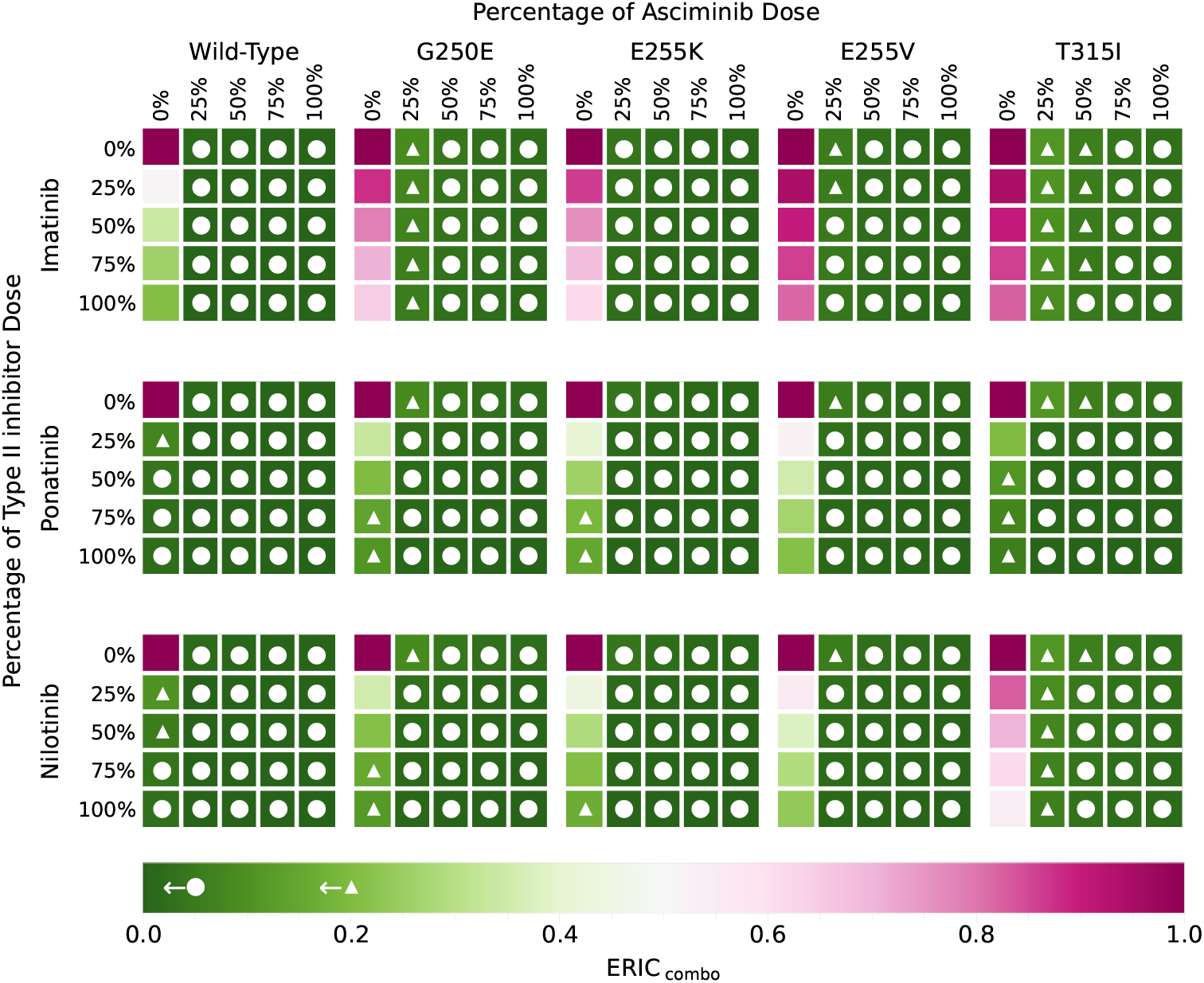
Heat map of the product of the Effective Ratios of IC_50_ (ERIC_combo_s) for each combination of drugs, dose and mutant with once-daily dose regime of asciminib. The inhibitor concentration values used are the midpoint of the range of inhibitor concentration on day 10. The white triangles and circles indicate ERIC that are under 0.2 and 0.05, respectively. An equivalent figure for twice-daily doses of asciminib can be found in the supporting information (Figure S6, with full ERIC_combo_ values detailed in Figures S7 to S9). Comparing this to Figure 6 shows how well ERIC_combo_ values can indicate effectiveness of treatment.

The success of combination therapy is measured in many ways. We will use zero interaction potency (ZIP) measurements and calculate the average difference between ZIP values and true behaviour (normally defined as *δ*) as a measure of synergy [26]. As we measure our model outputs using IRP, we define IRP_ZIP_ and *δ*_IRP_ to assess synergetic behaviour. ZIP scores and *δ*-values are normal measures of synergy, as such we do not fully define them here; however, further detail of our calculations for IRP_ZIP_ and *δ*_IRP_ can be found in section 4.2. A sample of synergy measurements can be seen in Figure 8 in comparison with IRP and the ERIC_combo_ values for 25% asciminib doses in the once-daily regime and 25% type II inhibitor doses. The equivalent figure for the twice-daily asciminib regime can be seen in the supporting information in Figure S10.

**Figure 8.**
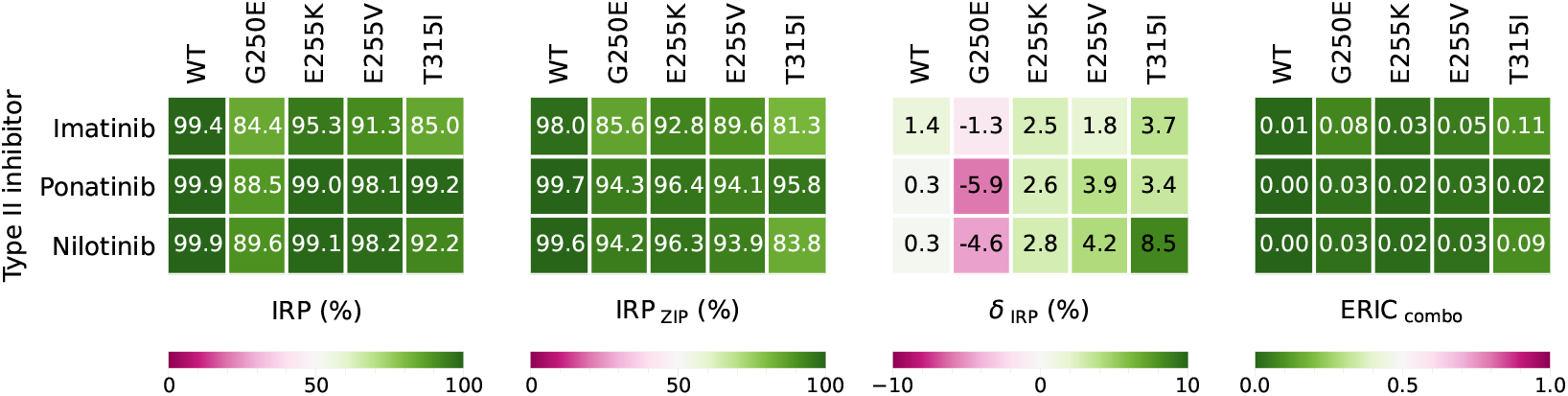
Comparison of IRP, IRP_ZIP_, *δ*_ZIP_, and the product of Effective Ratios of IC_50_ (ERIC_combo_) for 25% dose of type II inhibitors and 25% dose of once-daily asciminib. An equivalent figure for the twice-daily asciminib regime can be found in the supporting information (Figure S10, with full IRP_ZIP_ and *δ*_ZIP_ values detailed in Figures S11 to S16). The values of *δ*_IRP_ show low amounts of synergy for all mutants except G250E, which experiences low levels of antagonism, for these doses of combination therapy. The ERIC_combo_ values reflect the same trends seen in IRP and IRP_ZIP_.

The current mutation-drug matching guides of fold-IC_50_ values from the WT of each drug for each mutant can be seen in Figure 9. As we found no way to combine two fold-IC_50_ values to match the model’s outcomes, they have been left separated with the fold-IC_50_ value for asciminib in the upper right triangles and the fold-IC_50_ value for the type II inhibitors in the lower left triangles. As these values are independent of the inhibitor concentration, they cannot provide specific details in the same way ERIC_combo_ can, this is discussed further in the next section.

**Figure 9.**
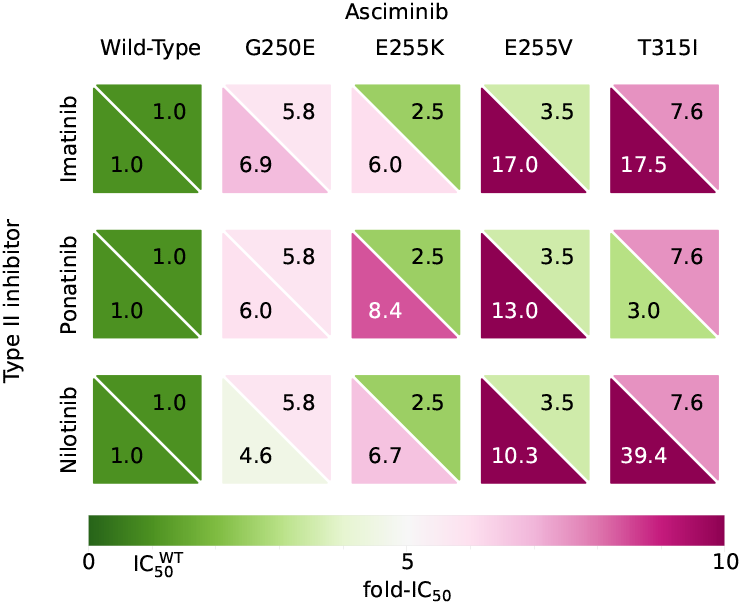
Fold-IC_50_ values from the WT of each drug for each mutant. Fold-IC_50_ value for asciminib in the upper right triangles and the fold-IC_50_ value for the type II inhibitors in the lower left triangles. Comparing to the IRP values from the simulations and ERIC_combo_ values calculated (Figures 6 and 7), we see no clear trend in fold-IC_50_ values and the behaviour of the system.

## 3 Discussion

There is a strong correlation between the amount of cobinding of the type II inhibitors and asciminib and the level of synergy in the system (see Figures 2 and 3). For example, the results show that G250E has much lower levels of cobinding and has antagonistic responses. This may be due to its lower Δ*G* value for the active to inactive state transition - i.e. the enzyme has a stronger preference of staying in the active state, which the type II inhibitors do not bind to. As in [13], we see that IRP gives a good representation of associated resistance and ERIC is a good indicator of expected IRP results (Figures 6 and 7). In this work, we have expanded on this by showing that as the ERIC values take the concentration of the inhibitors in patients into consideration and they can be combined through multiplication (ERIC_combo_) to give an indication of IRP values for combination treatment. This way of calculating ERIC_combo_ is based on an assumption that the cobinding of drugs is independent of the order of binding. The ERIC_combo_ values show a strong correlation to the IRP values that describe the system. As ERIC’s purpose is to help guide practitioners in treatment selection, overcomplicating the calculation with inclusion of extra details accounting for binding order, that we do not know the full details of, would not be beneficial. An alternative calculation for ERIC_combo_ for systems without cobinding (exclusive binding) is (compare to Equation 3):

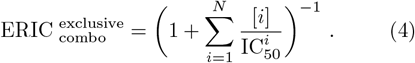

To examine the synergy of the system (as described in section 2) IRP_ZIP_ and *δ*_IRP_ values were calculated for the system. In Figure 8, the *δ*_IRP_ values show small levels of antagonism (*δ*_IRP_ < 0) for the G250E mutant and synergy for the other mutants (*δ*_IRP_ > 0). This may be linked to the lower levels of cobinding and G250E’s strong preference to remain in the active state. Experimental results at lower concentrations and other simulations show that synergy is expected for type II inhibitors and asciminib [18, 19, 20, 21, 22, 23]. This discrepancy could be due to differences in concentration or may be due to flaws in the calculation of the rates associated with the co-binding of asciminib and type II inhibitors.

These small amounts of synergy or antagonism allow for ERIC_combo_ to hold up well. Due to the way ERIC_combo_ is calculated, it gives an indication of IRP based on zero synergetic or antagonistic behaviour - which is also how ZIP values are calculated. Unlike ZIP values, no cell models or in vivo experiments are needed and the calculation process is much simpler. In cases where there is synergy, e.g. type II inhibitors with asciminib, ERIC_combo_ provides a good guide for treatment selection as it can provide a baseline for expected outcomes. However, with different inhibitors, like dasatinib, that interact antagonistically with asciminib [18], the ERIC_combo_ values will likely be an over estimation.

ERIC_combo_ provides a better indication of system behaviour than the traditional relative or fold-measures of IC_50_ values compared to the WT. Figure 9 shows the relative IC_50_ values for asciminib and type II inhibitors. Other than the general “less well than the WT” indication of these values (fold-IC_50_ *>* 1), we failed to find any way to combine two values to give an indication of behaviour in combination. Other methods for optimising drug combinations have been explored. For example, utilising large data bases of drug combination experiments to pinpoint optimal combination concentrations using *δ* values [26]; and using computational methods like machine learning to predict successful drug combinations [27]. Algorithms have also been designed specifically for optimising CML combination treatment using stochastic models based on cross-resistance properties and drug concentration [28]. These all need significantly more computational power than ERIC_combo_.

Furthermore, fold-IC_50_ values are independent of dose and are unable to inform dose selection in the way ERICs and ERIC_combo_s are. The opportunity to use ERIC_combo_ values to guide combination treatment could provide a wider array of treatment options for patients with high sensitivity to toxicity or resistant mutations. It may also extend into first-line treatment options for patients with blast- or accelerated-phase CML who generally require a more rapid and intense treatment.

The study of cancer is continuously broadening and bettering treatment options for patients; and ERIC and ERIC_combo_ could provide more accessible assessments for combination targeted and personalised therapies for other cancers.

## 4 Methods

The Abl1 enzyme is a key component of growth and multiplication signalling and the success of Abl1 inhibitors in the treatment of CML confirms the importance of reducing the product formation rate in the aim of achieving DMR in patients. This suggests (as our previous work shows [13]) that the relative change in product formation rate is a good indicator for resistance in therapy. This section outlines our methods for modelling the product rate and its response to treatments.

### 4.1 Modelling of drug resistant mutations

Following the methods in [13], the time-dependent concentration of each enzyme state can be written as:

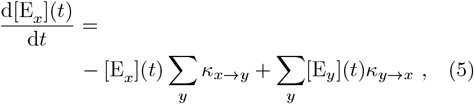

where *x* and *y* are arbitrary labels to represent two different enzyme states. The concentration of the enzyme in state E_*x*_ is denoted with [E_*x*_], and the rate constant for transition from state E_*x*_ to state E_*y*_ is given as *κ*_*x*→*y*_.

The key components of Equation 5 are the states the enzyme can be in and the rates of transition between these states. We outline the states for this simple model in section 4.1.1. Section 4.1.2 then examines the system and its behaviour in quasi-equilibrium conditions. This lays the foundation for deriving the relationships between the rate constants. If experimental rate constants are available from reliable kinetic experiments, these should be used. When such values are not known a priori, they are approximated as outlined in section 4.1.3.

In section 4.1.4, we outline the pharmacokinetics of asciminib and the type II inhibitors in the system. This will more accurately model the physiology of drug treatment, where the concentration fluctuates due to regular dosing at timed intervals. The last piece of the model is the initial conditions (section 4.1.5); these are found analytically, assuming a pre-existing quasiequilibrium steady state before treatment.

The time-dependent differential equations (as in Equation 5) were implemented using an Euler forward algorithm for the varying treatment concentrations with a time step of 1 ms. The mass balance equations for the model can be found in Table S1 in the supporting information.

#### 4.1.1 Determining the states of the system

The signalling of Abl1 is of high importance in the pathway of cell growth and proliferation and we focus on a measurement of the product formation rate of the phosphorylation of a substrate protein by Abl1 to indicate response to treatment. There are many combinations of active and inactive states of the Abl1 enzyme, substrate protein, ATP molecule, type II inhibitor, and asciminib to examine, but not all are relevant to the model. As in [13], we eliminate: the simultaneous binding of both type II inhibitor and ATP to Abl1; the substrate protein binding to the inactive Abl1; and the binding of the substrate protein or ATP alone (i.e. the binding of the ATP and the substrate protein is simplified into a single transition). The type II inhibitors that are examined bind only to the inactive state of the enzyme, so only the inactive state bound to a type II inhibitor needs to be considered. Lastly, once bound to the substrate protein and ATP molecule, the phosphorylation reaction proceeds rapidly. Therefore, we can also reject the combination of the substrate protein, ATP, and asciminib binding in combination.

This leaves a total of seven different states for the Abl1 enzymes in the system: unbound active enzyme, active enzyme bound to the ATP and substrate protein, active enzyme bound to asciminib, unbound inactive enzyme, inactive enzyme bound to a type II inhibitor, inactive enzyme bound to asciminib, and inactive enzyme bound to both asciminib and a type II inhibitor. These states can be seen in Figure 10. This figure also shows the transition rate constants between states, we discuss these in further detail in section 4.1.3.

**Figure 10.**
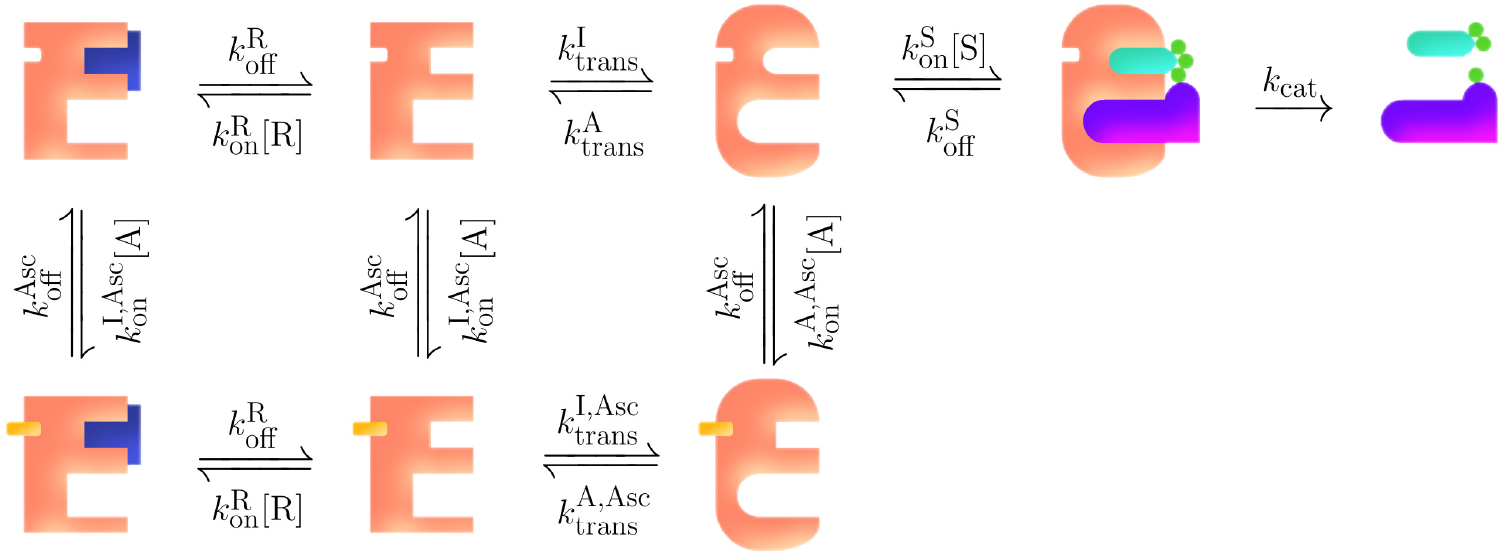
A simple illustration of the simplified model of the Abl1 enzyme system in focus. The states, molecules they bind to, the transitions between states, and the rate constants for each transition are shown. The Abl1 enzyme is represented by the peach-coloured E-shape in both its active (round corners) and inactive (sharp corners) states; a type II inhibitor is illustrated by the dark blue T-shape (either imatinib, ponatinib, or nilotinib); the small yellow shape represents an asciminib molecule; and the remaining shapes depict the ATP with its phosphate groups and the protein substrate that is phosphorylated. Transition arrows are accompanied by their rate constants. The rates between states will differ depending on the mutation and type II inhibitor present in the system.

#### 4.1.2 Fixed conditions and quasi-equilibrium

We first examine the system in a quasi-equilibrium with fixed inhibitor concentrations - i.e. a system with less complicated behaviour. This was used to identify the relationships between various rate constants using data from simulations and experiments.

We denote concentrations as [S] for the substrate, [R] for the type II inhibitors, and [A] for asciminib. The assumption for the concentration of ATP is that it is in surplus relative to the substrate and its concentration is therefore not treated explicitly by the model. Principles of chemical kinetics [29] and statistical mechanics provide tools to determine the proportion of the total enzyme concentration for each state considered in the model. These proportions are referred to as relative weights and are detailed in Figure 11. The preference of a mutation for the inactive or active state (i.e. whether the active or inactive state is more stable) is determined by the free-energy difference between these states. This is denoted as Δ*G*, which is defined by *ϵ*_A_ − *ϵ*_I_ where *ϵ*_A,I_ are the free energies of the active and inactive state of the enzyme, respectively.

**Figure 11.**
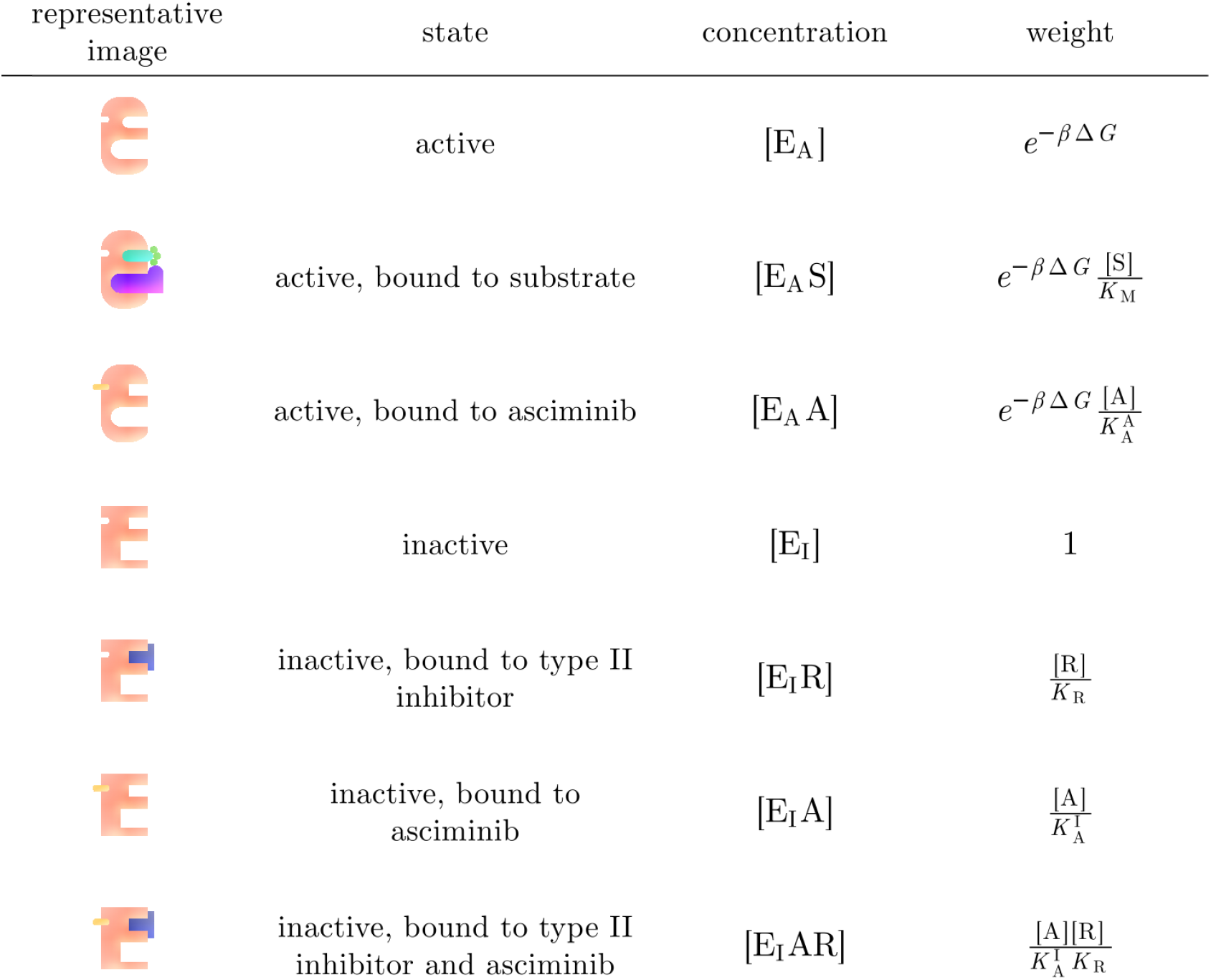
Details and labelling conventions for the enzyme states and their relative weights within the system in quasi-equilibrium. *β* is the reciprocal of the product of Boltzmann’s constant and the temperature (*β* = 1*/k*_*B*_*T*); Δ*G* is the change in Gibbs free energy between the active and inactive enzymes, and it describes the preference of the unbound enzyme between the active and inactive states; [S] is the concentration of the substrate (it is assumed that ATP is in surplus relative to the substrate and its concentration is therefore not treated explicitly by the model); *K*_M_ is the Michaelis constant of the binding of the substrate to the active enzyme state; [A] is the concentration of asciminib in the system; 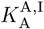 is the dissociation constant of the binding of asciminib for the enzyme in either active (A) or inactive (I) state; [R] is the concentration of the type II inhibitor in the system; and *K*_R_ is the dissociation constant of the binding of the type II inhibitor drug.

The product formation rate,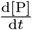, can be expressed in terms of *k*_cat_ (the turnover number or catalytic rate constant for the active enzyme) and the concentration of active enzyme bound to the substrate [E_A_S], as in [29]:

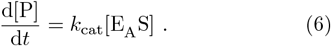

[E_A_S] can be expressed as a fraction of the total enzyme concentration, [E_tot_], in quasi-equilibrium. The weighting for a given state divided by the total of the weights for every state (*W*_tot_) gives the proportion of [E_tot_] that will be in that state. These weights are outlined in Figure 11. We can now rewrite the product formation rate as:

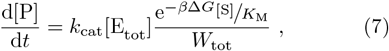

where

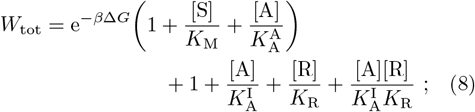

where *K*_M_ is the Michaelis’ constant of the binding of the substrate to the active enzyme state, 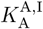 is the dissociation constant of the binding of asciminib for the enzyme in either active (A) or inactive (I) state, and *K*_R_ is the dissociation constant of the binding of the type II inhibitor drug.

#### 4.1.3 Quantitative determination of rate constants

Once we have an understanding of the product formation rate for a system in quasi-equilibrium, we proceed to capture how the product formation rate varies with fluctuating inhibitor concentrations. More details and calculations of the fluctuations in the inhibitor concentration can be found in section 4.1.4. To account for these dynamic changes, we must model the time-dependence of the concentrations of all the states and find the transition rates between them. In Equation 5, the rate constants are defined as (*κ*_*x*→*y*_ and *κ*_*y→x*_). Figure 10 depicts the rate constants we will calculate in the model. Table 2 bridges the two labelling conventions. Described below is our approach to obtaining these rate constants.

**Table 2:** To bridge two naming conventions (one more suitable to mathematical notation and one more conventional in enzymology), the table matches the rate constants as described in Equation 5 to those labelled in Figure 10.

| rate constant<br>of transitions<br>as in Equation 5 | rate constant<br>as labelled in<br>Figure 10 |
| --- | --- |
| $\kappa_{[E_I] \rightarrow [E_I R]}$ | $= k_{on}^R [R]$ |
| $\kappa_{[E_I R] \rightarrow [E_I]}$ | $= k_{off}^R$ |
| $\kappa_{[E_I] \rightarrow [E_I A]}$ | $= k_{on}^{I,Asc} [A]$ |
| $\kappa_{[E_A] \rightarrow [E_A A]}$ | $= k_{on}^{A,Asc} [A]$ |
| $\kappa_{[E_I A] \rightarrow [E_I]}$ | $= k_{off}^{Asc}$ |
| $\kappa_{[E_I R] \rightarrow [E_I A R]}$ | $= k_{on}^{I,Asc} [A]$ |
| $\kappa_{[E_I A R] \rightarrow [E_I R]}$ | $= k_{off}^{Asc}$ |
| $\kappa_{[E_I A] \rightarrow [E_I A R]}$ | $= k_{on}^R [R]$ |
| $\kappa_{[E_I A R] \rightarrow [E_I A]}$ | $= k_{off}^R$ |
| $\kappa_{[E_I] \rightarrow [E_A]}$ | $= k_{trans}^I$ |
| $\kappa_{[E_A] \rightarrow [E_I]}$ | $= k_{trans}^A$ |
| $\kappa_{[E_I A] \rightarrow [E_A A]}$ | $= k_{trans}^{I,Asc}$ |
| $\kappa_{[E_A A] \rightarrow [E_I A]}$ | $= k_{trans}^{A,Asc}$ |
| $\kappa_{[E_A] \rightarrow [E_A S]}$ | $= k_{on}^S [S]$ |
| $\kappa_{[E_A S] \rightarrow [E_A]}$ | $= k_{off}^S$ |
| $\kappa_{[E_A S] \rightarrow [E_A] + Products}$ | $= k_{cat}$ |

##### Abl1 type II inhibitor binding rate constants 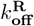 and 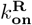

The dissociation constant *K*_R_ for type II inhibitors can be written as [13]:

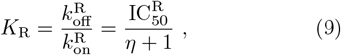

where *η* is used as a shorthand and is defined as

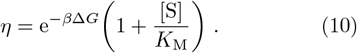

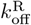 is found using measured residence times *t*_R_ as:

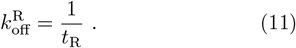

The residence times and their associated 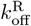 values can be found in Table 3. IC_50_ values can be found in Table 4.

**Table 3:** Residence times, *t*_R_, of the inhibitor molecules unbinding from the wild-type from [30] and the rates of 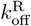 calculated from them.

| ABL1<br>inhibitor | $t_R$<br>(min) | $k_{off}^R$<br>($\text{min}^{-1}$) |
| --- | --- | --- |
| imatinib | 17 | 0.059 |
| ponatinib | 205 | 0.00488 |
| nilotinib | 50 | 0.020 |

**Table 4:** Δ*G* values are from [13]. The values for *k*_cat_ and *K*_M_ are sourced from [31]. All the IC_50_ values for the type II inhibitors (imatinib, ponatinib and nilotinib) are from [32]. The IC_50_ values for asciminib are taken from [33].

| Mutation | $\Delta G$<br>(kcal mol<sup>-1</sup>) | $k_{\text{cat}}$<br>(min<sup>-1</sup>) | $K_M$<br>($\mu\text{M}$) | IC<sub>50</sub>(nM) | | | |
| --- | --- | --- | --- | --- | --- | --- | --- |
|  |  |  |  | Imatinib | Ponatinib | Nilotinib | Asciminib |
| Wild-type | -1 | 66 | 17 | 527.0 | 2.1 | 17.7 | 3.8 |
| G250E | -3.2 | 175.2 | 14.3 | 3613.0 | 12.5 | 80.7 | 22 |
| E255K | -1.5 | 63.6 | 15.6 | 3174 | 17.6 | 118.4 | 9.5 |
| E255V | -1.4 | 6.8 | 22.1 | 8953 | 27.2 | 182.3 | 13.3 |
| T315I | -0.9 | 12.2 | 7.2 | 9221 | 6.3 | 697.1 | 28.7 |

Equations 9 to 11 can be used to find the 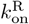 values for the WT for each type II inhibitor. Once bound, a type II inhibitor and a resistance associated mutants of a kinase behave in a similar manner to the inhibitor in combination with the wild-type enzyme [34]. We use this idea to set the 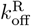 for these type II inhibitors (imatinib, ponatinib and nilotinib), as the same across the WT and the mutants. The values for the *K*_R_ and 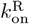 can be found in Table 5.

**Table 5:** Type II inhibitor dissociation constants and binding rates. Calculated using Equation 9.

| Mutation | $K_R$(nM) | | | $k_{\text{on}}^R (\times 10^{-3} \text{nM}^{-1} \text{min}^{-1})$ | | |
| --- | --- | --- | --- | --- | --- | --- |
|  | imatinib | ponatinib | nilotinib | imatinib | ponatinib | nilotinib |
| Wild-type | 203.6 | 0.811 | 13.48 | 6.01 | 0.290 | 1.48 |
| G250E | 1338 | 4.63 | 6.25 | 1.05 | 0.0441 | 3.20 |
| E255K | 12021 | 6.66 | 68.49 | 0.732 | 0.0491 | 0.292 |
| E255V | 3651 | 11.1 | 117.7 | 0.440 | 0.0162 | 0.170 |
| T315I | 2721 | 1.86 | 497.7 | 2.63 | 0.0217 | 0.0402 |

##### Asciminib binding rate constants: 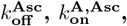 and 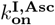

The value of 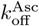 was estimated by performing simulations with different values to examine its effect on the response of the system. The values tested were of the same order of magnitude as those of 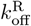 outlined in Table 3. Further details and results of these tests can be found in the supporting information (Figure S17). Across all mutations, the unbinding rate for both the active and inactive state is selected to be 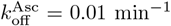. In order to find the binding rates for asciminib in the active and inactive states, we must first derive the dissociation constants for those states.

To find these values we look to the behaviour of the system with asciminib and without type II inhibitors present and compare that to the system with no inhibitors present. Without any inhibitors, we can rewrite Equation 7 as

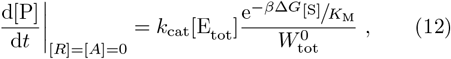

where

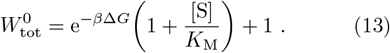

When [R]=0 and 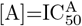, we can write the product rate as

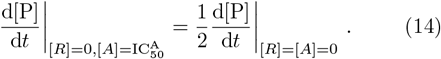

Substituting [R]=0 and 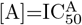 into Equation 8, we can form an expression for the relationship between 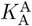 and 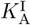:

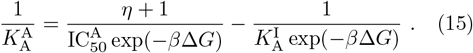

For the sake of simplicity we define another relationship between the two dissociation constants as 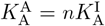 and form expressions for 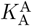 and 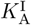 in terms of *n*.

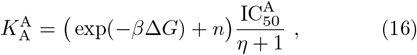

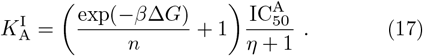

We can see that 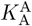 is linear with *n* and the relationship between 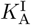 and *n* is non-linear and is limited by *n >* 0 and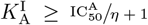 .The binding of asciminib to the inactive state is more favourable than its binding to the active state of the WT enzyme [18], therefore we apply the bound of 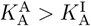. This limits the value of of the inactive dissociation constant to be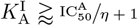. We chose to select 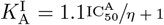, which gives

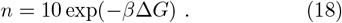

Using the relationship between the dissociation constant and the binding and unbinding rates (as in Equation 9) the binding constants for the active and inactive states can be found. IC_50_ values can be found in Table 4 and the values for 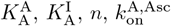, and 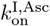 can be found in Table 6.

**Table 6:** Asciminib dissociation constants and binding rates. Calculated using Equations 16 to 18.

| Mutation | $n$ | $K_A^A$<br>(nM) | $K_A^I$<br>(nM) | $k_{\text{on}}^{A, \text{Asc}}$<br>($\times 10^{-3} \text{nM}^{-1} \text{min}^{-1}$) | $k_{\text{on}}^{I, \text{Asc}}$<br>($\times 10^{-3} \text{nM}^{-1} \text{min}^{-1}$) |
| --- | --- | --- | --- | --- | --- |
| Wild-type | 1.97 | 6.28 | 3.18 | 1.59 | 3.14 |
| G250E | 70.1 | 131.4 | 1.87 | 0.0761 | 5.34 |
| E255K | 4.44 | 26.8 | 6.04 | 0.373 | 1.65 |
| E255V | 3.78 | 35.7 | 9.45 | 0.280 | 1.06 |
| T315I | 1.68 | 37.8 | 22.5 | 0.265 | 0.444 |

##### Substrate binding: 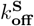 and 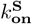

The rate constants associated with substrate binding are estimated using experimental results (specifically values for *k*_cat_ and *K*_M_) and an examination of the behaviour of the WT. The relationship between the substrate binding and unbinding rates, *k*_cat_ and *K*_M_ is:

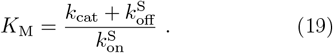

The values for *K*_M_ and *k*_cat_ can be found in Table 4. As with [13], we use a simple estimation that, for the WT 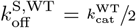 The value for 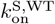 for the WT can then be calculated using Equation 19. For the substrate binding rates for mutants, we employ the same algorithm as outlined in [13]: 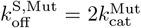 if the resulting 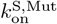 (using Equation 19) is less than 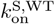, we assign 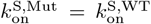 and a new 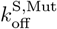 is calculated using Equation 19). The final values of these estimates are shown in Table 7.

**Table 7:** The substrate binding and unbinding rates for the WT and each mutant Abl1 enzyme.

| Mutation | $k_{\text{off}}^S$<br>($\text{min}^{-1}$) | $k_{\text{on}}^S$<br>($\mu\text{M}^{-1}\text{min}^{-1}$) |
| --- | --- | --- |
| Wild-type | 33.0 | 5.82 |
| G250E | 350 | 36.8 |
| E255K | 127 | 12.2 |
| E255V | 122 | 5.82 |
| T315I | 29.7 | 5.82 |

##### Transitions between the active and inactive enzyme states: 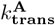 and 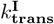

The relationship between the transition rates is

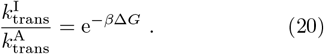

The values for Δ*G* are taken from results produced by free energy perturbation simulations in [13] and can be found in Table 4. As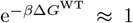, we assume that 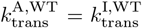. We find this also applicable for majority of the mutants, with G250E having a larger Δ*G*, so has a smaller approximated transition rate. We have approximated that 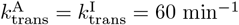 for the WT and mutants E255K, E255V and T315I. For G250E, 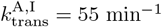.

##### Transitions between the active and inactive enzyme states while bound to asciminib: 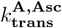 and 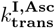

The free energies of the cobinding of nilotinib and asciminib have been simulated and measured [18]. This includes information about the transition between the active and inactive states while bound to asciminib. We use these values in conjunction with the 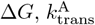 and 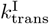 values from [13] (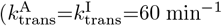 for WT and mutants, with the exception of G250E, where this value is 55 min^*−*1^) to find 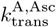 and 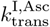 for each mutation.

We construct the following closed thermodynamic cycle:

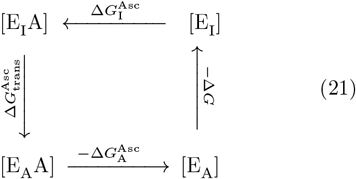

where 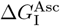 is the change in free energy for asciminib binding to the inactive state,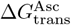 is the change in free energy from the inactive to the active state while bound to asciminib, and 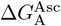 is the change in free energy for the binding of asciminib to the active state. From [18], we obtain 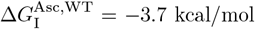 and 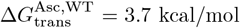. As Δ*G*^WT^ = −1 kcal/mol (as in Table 4), we find 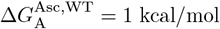. Generally, 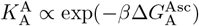 so by comparing the 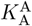 values for the WT and each mutation, we can write:

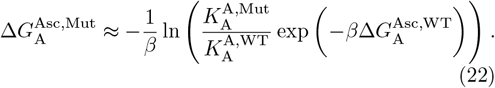

With the same treatment for the inactive state’s dissociation constant for asciminib, we form:

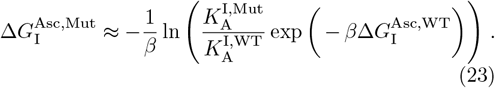

This allows us to calculate 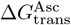 using:

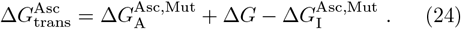

Using the assumption that the change in the rate of transition from the active state to the inactive state is of a similar size for each mutation and the WT (i.e. make 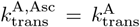), then we can readily calculate 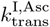 using:

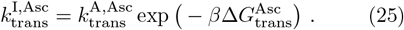

The values for 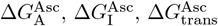, and 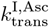 can be found in Table 8.

**Table 8:** Values for 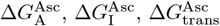, and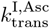.

| Mutation | $\Delta G_{\text{A}}^{\text{Asc}}$<br>(kcal mol<sup>-1</sup>) | $\Delta G_{\text{I}}^{\text{Asc}}$<br>(kcal mol<sup>-1</sup>) | $\Delta G_{\text{trans}}^{\text{Asc}}$<br>(kcal mol<sup>-1</sup>) | $k_{\text{trans}}^{\text{I,Asc}}$<br>(min<sup>-1</sup>) |
| --- | --- | --- | --- | --- |
| Wild-type | 1 | -3.7 | 3.7 | 0.148 |
| G250E | -0.110 | -2.61 | -0.7 | 171.3 |
| E255K | 0.416 | -3.78 | 2.7 | 0.749 |
| E255V | 0.143 | -4.16 | 2.9 | 0.542 |
| T315I | -0.00908 | -4.81 | 3.9 | 0.107 |

#### 4.1.4 Time-dependent drug concentrations in patients

There has been some investigation into how combination treatment affects inhibitor concentrations in patients [35]; however, the changes are small. Therefore, we treat the concentrations of asciminib and the type II inhibitor in the combination as independent of each other. To model realistic conditions, we construct a varying concentration of the inhibitors driven by multiple consistently spaced doses of inhibitor - i.e. like a patient taking medication daily or twice-daily.

Each dose taken causes an increases in concentration as the drug is absorbed through digestion. After some time, the effect of eliminating the drug overcomes absorption, and there is a decrease in concentration. Any remaining concentration of the drug at the time of the next dose is included in the total concentration with the next dose. Over time this effect can build up to a consistent baseline amount and the concentration of drug enters a steady state, fluctuating between a constant maximum and minimum.

We calculate the plasma concentration of the inhibitor *C*(*t*) at time *t* from the first dose of inhibitor with the following pharmacokinetic equation [36]

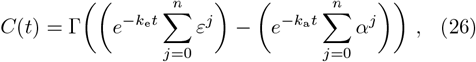

where *n* is the number of doses (12 or 24 hour periods depending on the inhibitor) since the first dose. *k*_*a*_ and *k*_*e*_ are the absorption and elimination rate constants of the Abl1 inhibitor. Γ, *α*, and *ε* are constants that are given by:

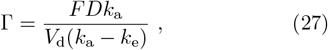

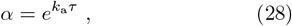

and

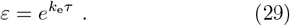

*F* is the bioavailability, *D* is the daily dose taken in moles, *V*_*d*_ is the volume of distribution, and *τ* is the dosing interval. The bioavailability of imatinib is 98% [37], but less precise values or qualitative descriptions are found for the other inhibitors. The bioavailability of ponatinib and asciminib were described as “high” [38] and “orally bioavailable” [39], respectively. Therefore, the value for *F* was assumed to be 1 for these three inhibitors. The bioavailibility for nilotinib was found as “moderate (17-44%)” and “predicted to be low (< 25%)” [40], so for nilotinib, we have selected *F* = 0.25.

The elimination rate, *k*_e_, was calculated using half-life values, 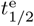, as 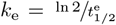. Usually, medicinal doses are provided as the mass of the active ingredient, hence the molar mass was used to calculate the dose in moles, *D*. The medicinal doses we use as baselines in this work are imatinib daily doses of 400 mg, ponatinib daily doses of 45 mg, nilotinib twice-daily doses of 400 mg, asciminib daily doses of 80 mg, and asciminib twice-daily doses of 40 mg. We also modelled 25, 50 and 75% doses of these medications as tablet-cutters are widely available, whereas varying doses may not be. We chose a system with regular doses of treatment, hence the time between doses, *τ*, is 12 hours for twice-daily treatments and 24 hours for daily treatments. Values for the pharmokinetic calculations can be found in Table 9.

**Table 9:** Values to model the time-dependent drug concentrations in the system based on observed data found and derived from a variety of sources. Sources listed respective to imatinib, ponatinib, nilotinib and asciminib. Dosing intervals, *τ*, were found in [41, 42, 43] and [39]. Absorption constants, *k*_a_ were taken from [44, 42, 45] and [46]. The elimination half-lifes,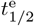, that the elimination constants, *k*_e_, were calculated from are from [47, 48, 43], and [39]. For the dose, *D*, we chose masses described as daily doses in”typical treatment” [41, 42, 43] and [39]. The volumes of distribution, *V*_d_, were taken from [47, 49, 43], and [46]. Values for Γ, *α* and *ε* were calculated using Equations 27-29.

|  | Type II inhibitor |  |  | Asciminib regime |  |
| --- | --- | --- | --- | --- | --- |
|  | imatinib | ponatinib | nilotinib | once-daily | twice-daily |
| $\tau$ (h) | 24 | 24 | 12 | 24 | 12 |
| $k_a$ (h<sup>-1</sup>) | 0.94 | 1.302 | 0.3 | 1.96 | |
| $t_{1/2}^e$ (h) | 18 | 24 | 16 | 8 | |
| $k_e$ (h<sup>-1</sup>) | 0.0385 | 0.0289 | 0.0433 | 0.0866 | |
| $D$ (mg) | 400 | 45 | 400 | 80 | 40 |
| $M$ (g/mol) | 493.603 | 532.6 | 529.5 | 449.8 | |
| $D$ (μmol) | 810.4 | 84.49 | 755.4 | 177.8 | 88.92 |
| $V_d$ (L) | 435 | 1223 | 579 | 111 | |
| $\Gamma$ (nM) | 1942.5 | 70.65 | 381.21 | 1676.3 | 838.14 |
| $\alpha$ | 6.28E+09 | 3.72E+13 | 3.66E+01 | 2.69E+20 | 1.64E+10 |
| $\varepsilon$ | 2.52 | 2 | 1.68 | 8.00 | 2.83 |

#### 4.1.5 Statistical mechanics of the initial conditions of the system without treatment

The initial conditions for the the WT and each mutant in this combination treatment model do not differ from those in [13], as there is no inhibitor present. We again rewrite Equation 7 under this condition and get equations for the initial inhibitor-free states:

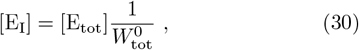

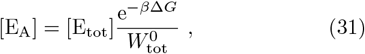

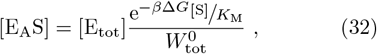

where,

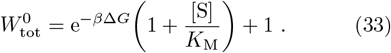

The total concentration of enzymes [E_tot_] is set to 1 *µ*M [50, 51].

### 4.2 Calculating synergy

Calculating the synergy for this system follows similar methods to those outlined in [26] for the calculation of ZIP and *δ* values. As our system is not subject to random variation as in vitro experiments may be, we have no need to fit data to curves and we simplify the method. For the ZIP values, we calculate a combined IRP (IRP_ZIP_) that assumes that a drug’s effects are not affected by the presence of a second drug and vice versa. We write the IRP_ZIP_ in terms of the IRPs for a type II inhibitor at concentration [R] and asciminib at concentration [A]:

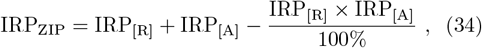

where IRP_[R]_ is the IRP of the system when only that concentration of the type II inhibitor is present and IRP_[A]_ is the IRP for the system with only that concentration of asciminib present. The difference between this IRP_ZIP_ value and the true IRP from the simulation for each combination of concentrations [R] of the type II inhibitor and [A] of asciminib gives a measure of synergy:

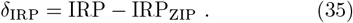

