## Supporting Information for "A computational dynamic model of combination treatment for type II inhibitors with asciminib"

February 21, 2025

#### Mass balance equations

| change | | calculation |
|---|---|---|
| $\Delta[\mathrm{E_A}]$ | $=$ | $\big((k_{\mathrm{off}}^{\mathrm{S}} + k_{\mathrm{cat}})[\mathrm{E_A S}] - k_{\mathrm{on}}^{\mathrm{S}}[\mathrm{S}][\mathrm{E_A}] + k_{\mathrm{trans}}^{\mathrm{I}}[\mathrm{E_I}] - k_{\mathrm{trans}}^{\mathrm{A}}[\mathrm{E_A}]$ |
| | | $+k_{\mathrm{off}}^{Asc}[\mathrm{E_A A}] - k_{\mathrm{on}}^{A,Asc}[\mathrm{A}][\mathrm{E_A}]\big)\Delta t$ |
| $\Delta[\mathrm{E_A S}]$ | $=$ | $\big(k_{\mathrm{on}}^{\mathrm{S}}[\mathrm{S}][\mathrm{E_A}] - (k_{\mathrm{off}}^{\mathrm{S}} + k_{\mathrm{cat}})[\mathrm{E_A S}]\big)\Delta t$ |
| $\Delta[\mathrm{E_A A}]$ | $=$ | $\big(k_{\mathrm{on}}^{A,Asc}[\mathrm{A}][\mathrm{E_A}] - k_{\mathrm{off}}^{Asc}[\mathrm{E_A A}] + k_{\mathrm{trans}}^{\mathrm{I,Asc}}[\mathrm{E_I A}] - k_{\mathrm{trans}}^{\mathrm{A,Asc}}[\mathrm{E_A A}]\big)\Delta t$ |
| $\Delta[\mathrm{E_I}]$ | $=$ | $\big(k_{\mathrm{trans}}^{\mathrm{A}}[\mathrm{E_A}] - k_{\mathrm{trans}}^{\mathrm{I}}[\mathrm{E_I}] + k_{\mathrm{off}}^{\mathrm{R}}[\mathrm{E_I R}] - k_{\mathrm{on}}^{\mathrm{R}}[\mathrm{R}][\mathrm{E_I}] + k_{\mathrm{off}}^{Asc}[\mathrm{E_I A}] - k_{\mathrm{on}}^{I,Asc}[\mathrm{A}][\mathrm{E_I}]\big)\Delta t$ |
| $\Delta[\mathrm{E_I R}]$ | $=$ | $\big(k_{\mathrm{on}}^{\mathrm{R}}[\mathrm{R}][\mathrm{E_I}] - k_{\mathrm{off}}^{\mathrm{R}}[\mathrm{E_I R}] + k_{\mathrm{off}}^{\mathrm{Asc}}[\mathrm{E_I AR}] - k_{\mathrm{on}}^{\mathrm{I,Asc}}[\mathrm{A}][\mathrm{E_I R}]\big)\Delta t$ |
| $\Delta[\mathrm{E_I A}]$ | $=$ | $\big(k_{\mathrm{on}}^{\mathrm{I,Asc}}[\mathrm{A}][\mathrm{E_I}] - k_{\mathrm{off}}^{\mathrm{Asc}}[\mathrm{E_I A}] + k_{\mathrm{off}}^{\mathrm{R}}[\mathrm{E_I AR}] - k_{\mathrm{on}}^{\mathrm{R}}[\mathrm{R}][\mathrm{E_I A}]\big)\Delta t$ |
| $\Delta[\mathrm{E_I AR}]$ | $=$ | $\big(k_{\mathrm{on}}^{\mathrm{R}}[\mathrm{R}][\mathrm{E_I A}] - k_{\mathrm{off}}^{\mathrm{R}}[\mathrm{E_I AR}] + k_{\mathrm{on}}^{\mathrm{I,Asc}}[\mathrm{A}][\mathrm{E_I R}] - k_{\mathrm{off}}^{\mathrm{A}}[\mathrm{E_I AR}]\big)\Delta t$ |
| $\Delta[\mathrm{E_{tot}}]$ | $=$ | $\Delta[\mathrm{E_A}] + \Delta[\mathrm{E_A S}] + \Delta[\mathrm{E_A A}] + \Delta[\mathrm{E_A R}] + \Delta[\mathrm{E_I}] + \Delta[\mathrm{E_I A}] + \Delta[\mathrm{E_I AR}]$ |
| | $=$ | $0$ |

Table S1: Table outlining the equations for the change in population of each state in the system using Euler integration of Equation 5 over time step $\Delta t$. The mass balance equation for the change in the total concentration, $\Delta[\mathrm{E_{tot}}]$, is given and shown to be zero.

### Sample of enzyme state outputs

To mirror Figure 2 in the main text, which shows the effect on the states of the T315I enzyme of 50% doses of imatinib and once-daily asciminib, here we include two further examples (Figure S1).

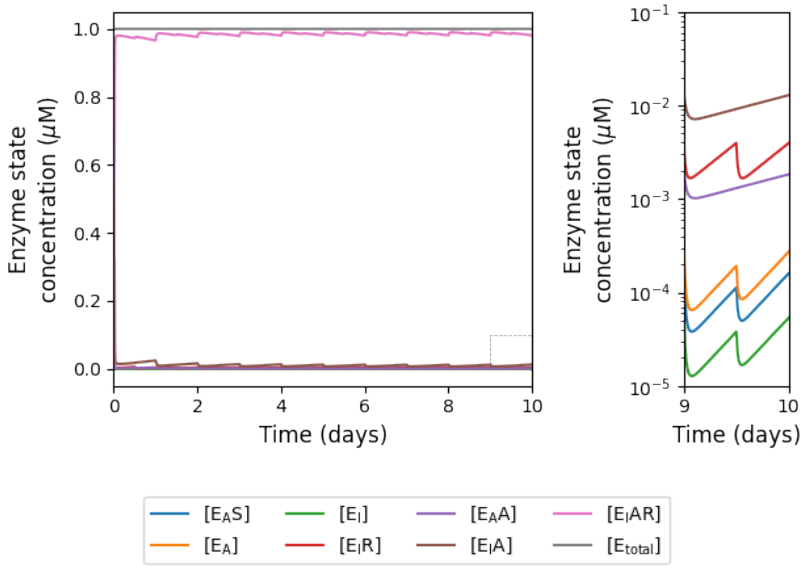


(a) WT with 25% ponatinib and 25% twice-daily asciminib.

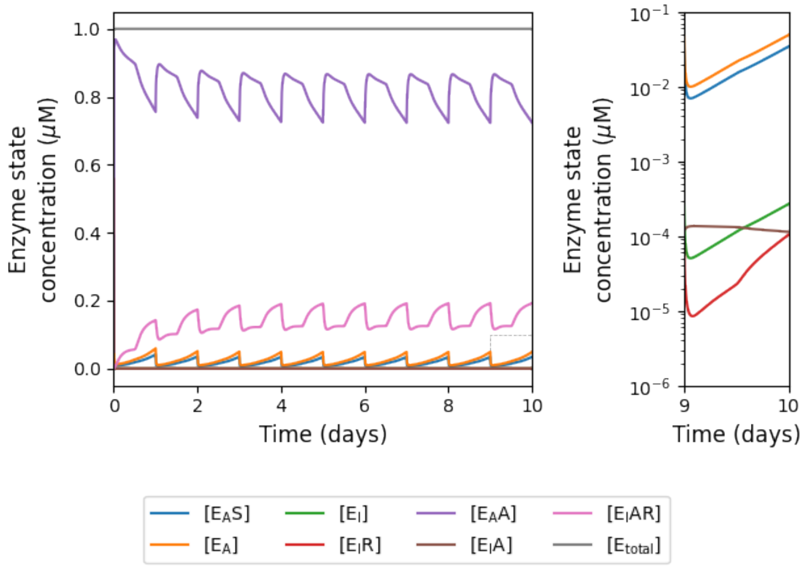


(b) G250E with 25% nilotinib and 75% once-daily asciminib.

Figure S1: A sample of the effect on enzyme states under different doses for different mutations.

### Inhibitory reduction prowess (IRP) overview - twice daily asciminib regime

In the main text we present an overview of our IRP results for once-daily asciminib doses for our model (Figure 6), here we show the equivalent results for twice-daily asciminib doses.

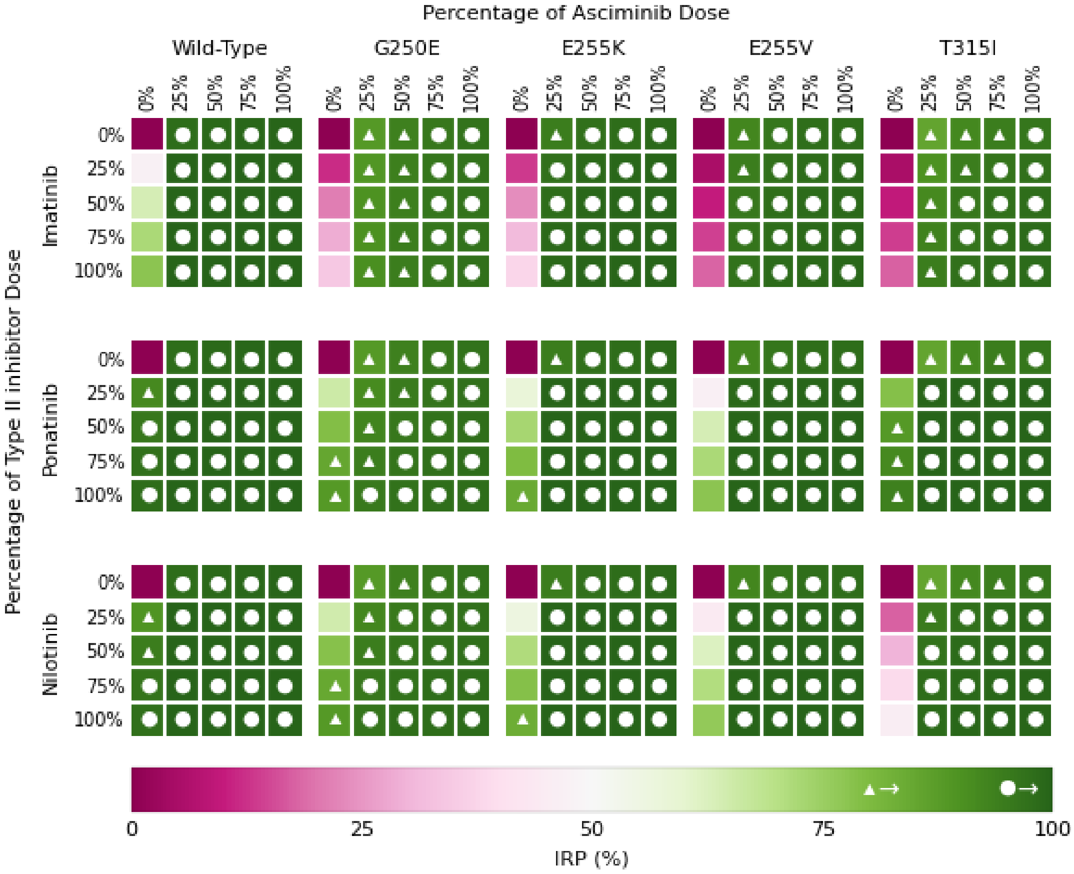


Figure S2: Heat map of IRP values for the midpoint of the range of product formation rate on day 10 for different combinations of doses of type II inhibitor and twice-daily doses of asciminib. The white triangles and circles indicate IRPs that are over 80% and 95%, respectively. An equivalent figure for once-daily doses of asciminib can be found in the main text in Figure 6.

### Full inhibitory reduction prowess (IRP) heat maps

In the main text in Figure 6 and in supporting information Figure S2, we present an overview of our IRP results, here we show the values calculated for every combination of dose, mutation, and inhibitor.

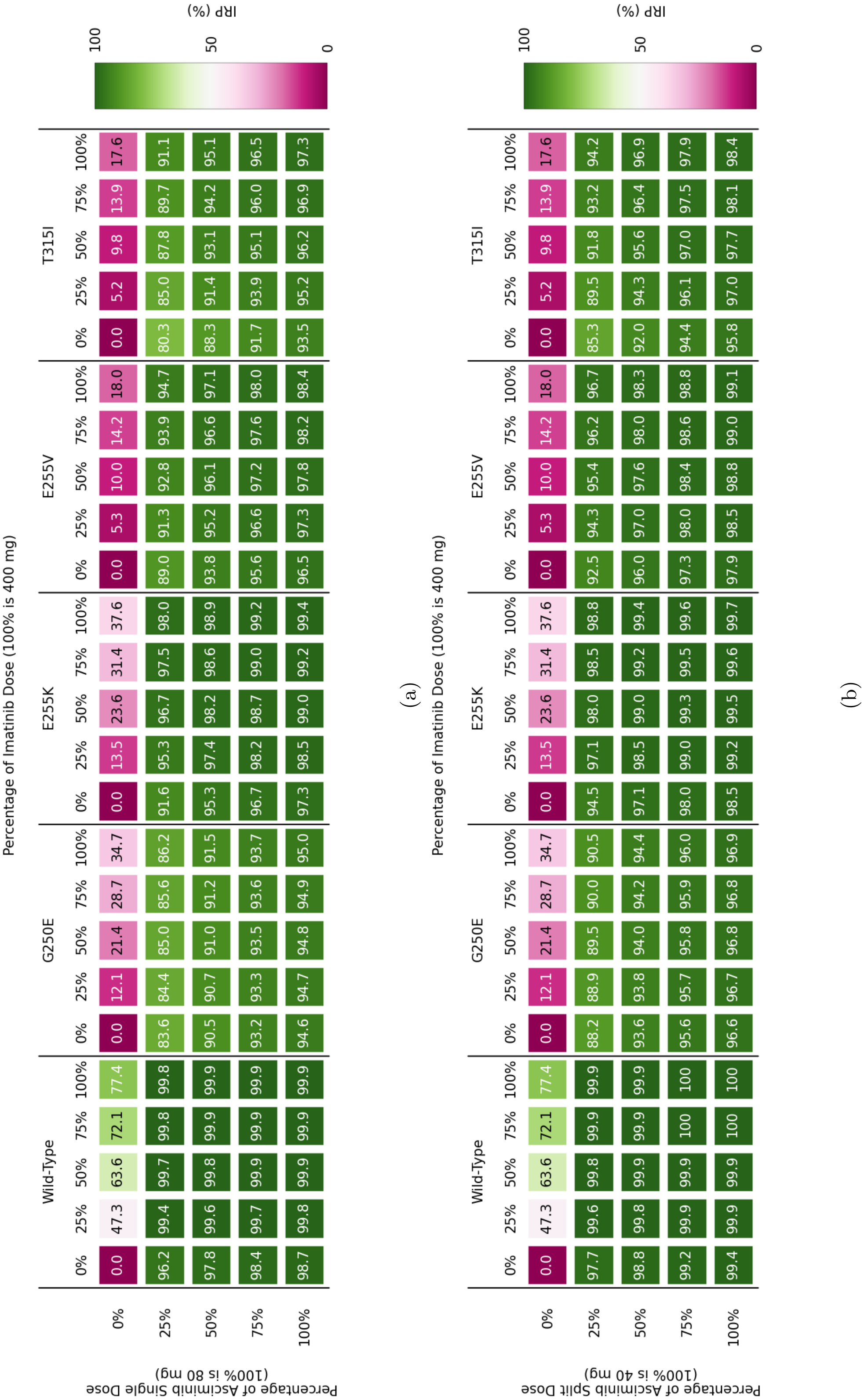


Figure S3: IRP full details of combinations of mutations and doses for imatinib with (a) once-daily and (b) twice-daily asciminib.

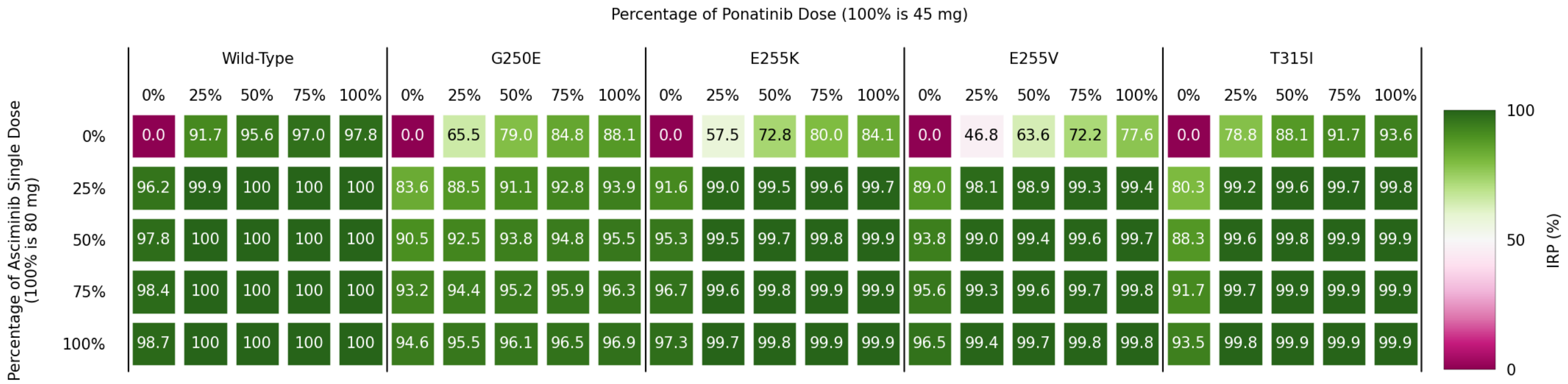


(a)

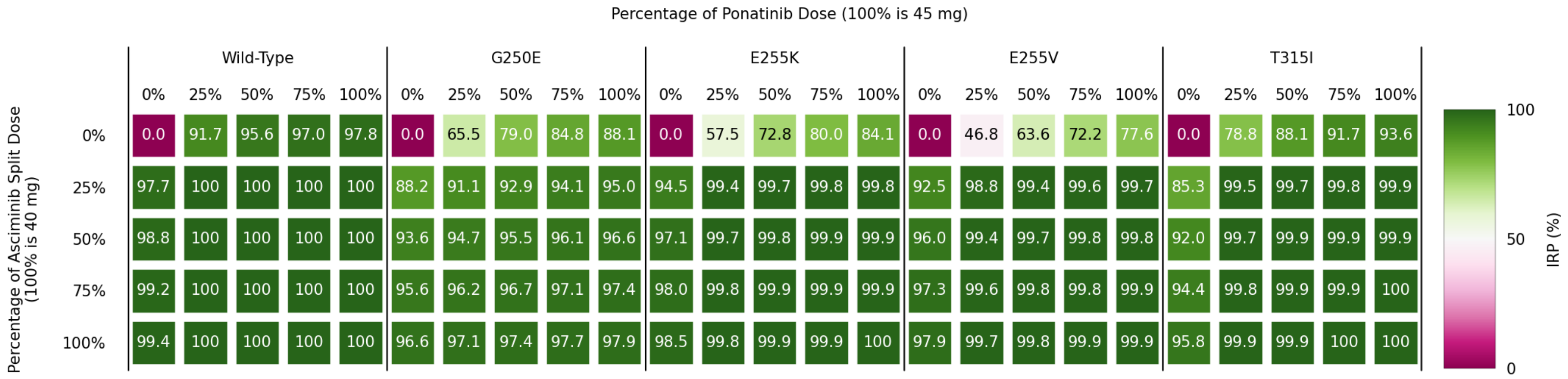


(b)

Figure S4: IRP full details of combinations of mutations and doses for ponatinib with (a) once-daily and (b) twice-daily asciminib.

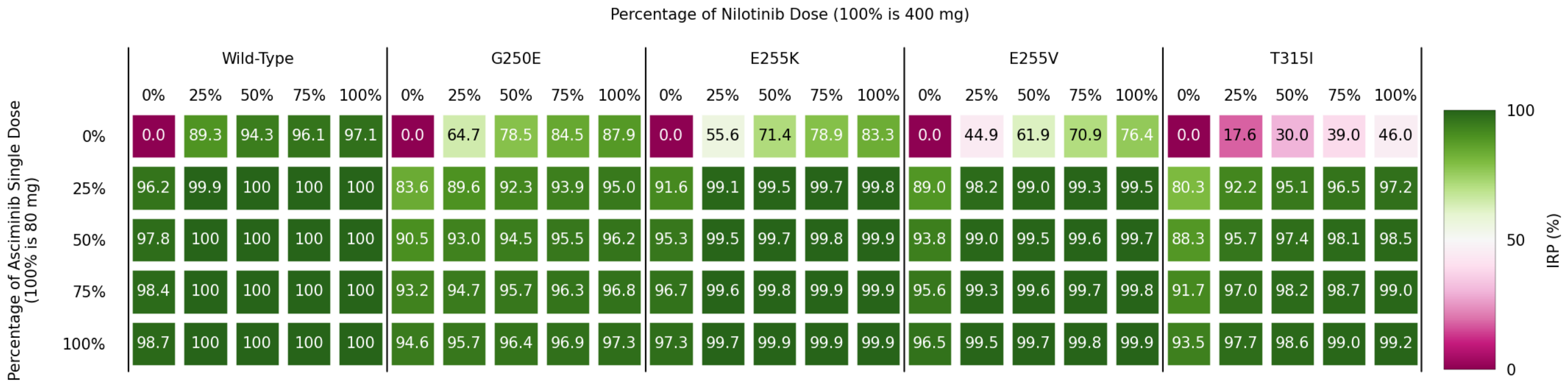


(a)

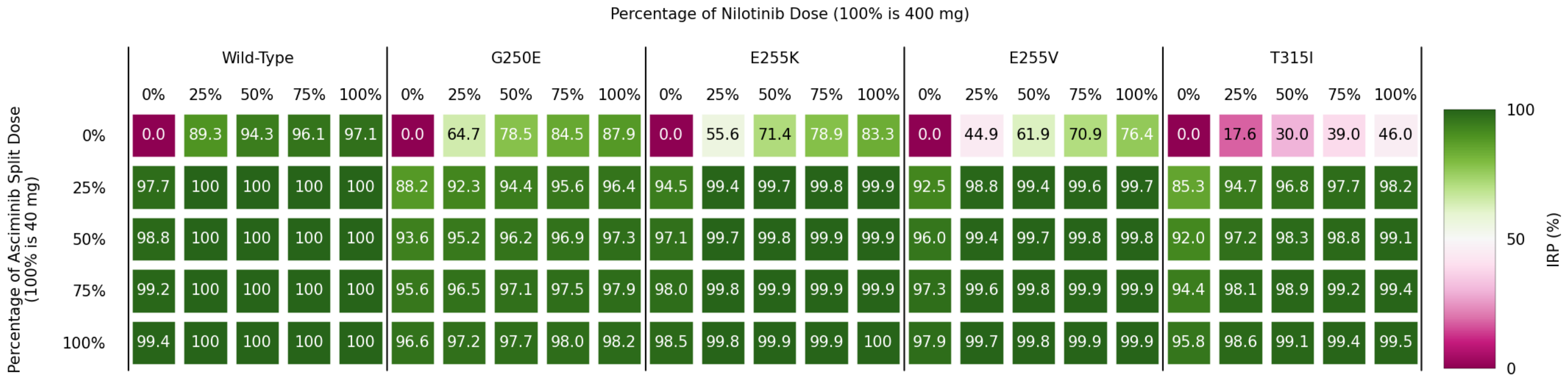


(b)

Figure S5: IRP full details of combinations of mutations and doses for nilotinib with (a) once-daily and (b) twice-daily asciminib.

#### Effective ratio of $IC_{50}$ in combination overview - twice-daily asciminib regime

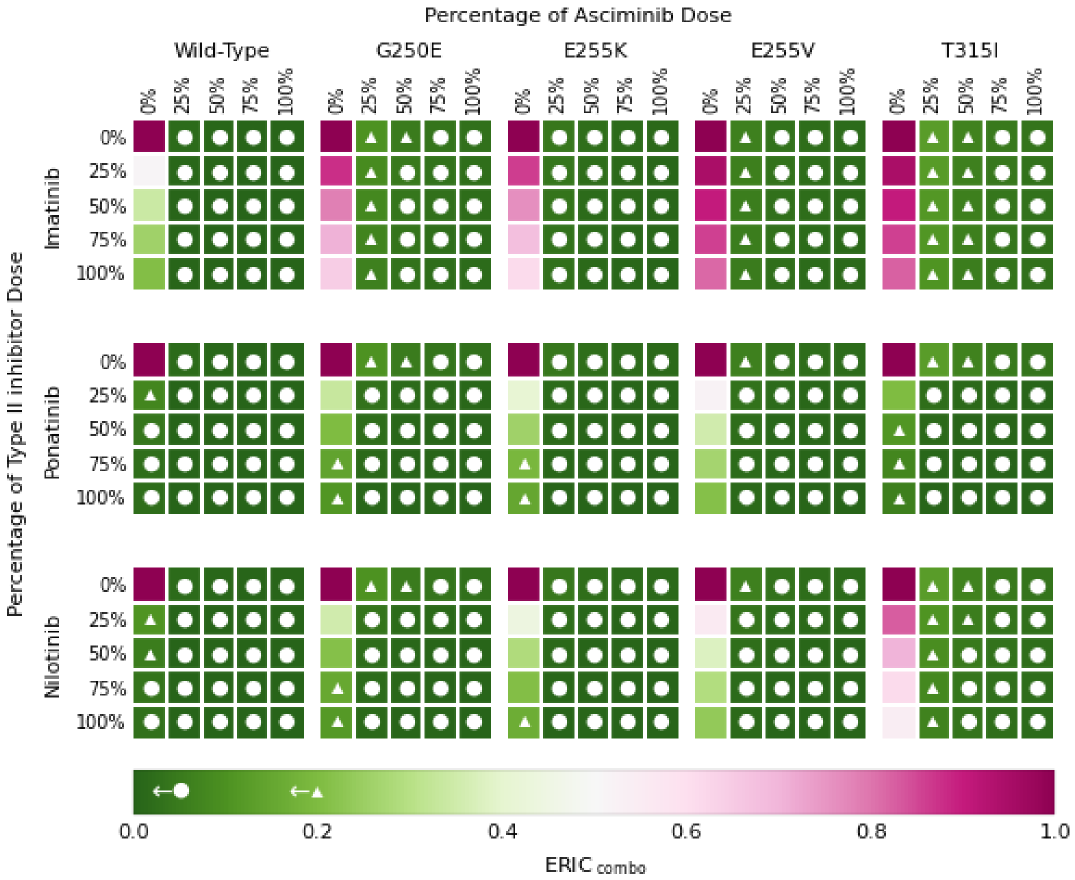


Figure S6: Heat map of the product of the Effective Ratios of $IC_{50}$ ($ERIC_{combo}$s) for each combination of drugs, dose and mutant with twice-daily dose regime of asciminib. The white triangles and circles indicate ERIC that are under 0.2 and 0.05, respectively. An equivalent figure for once-daily doses of asciminib can be found in Figure 7 in the main text.

### Full Effective Ratios of $IC_{50}$ ($ERIC_{combo}$s) heat maps

In the main text in Figure 7 and in supporting information Figure S6, we present an overview of our $ERIC_{combo}$ values, here we show the values calculated for every combination of dose, mutation, and inhibitor.

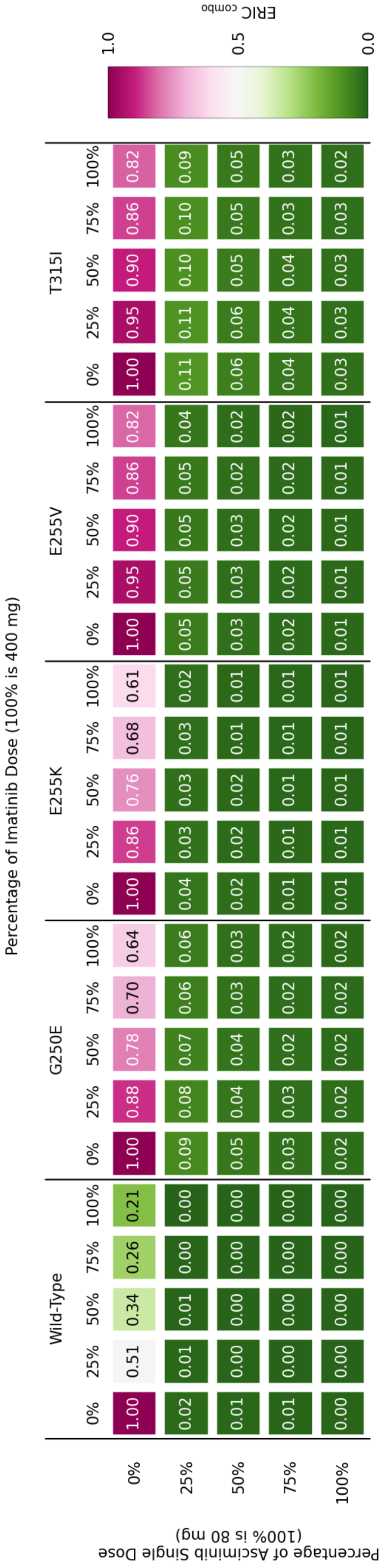


(a)

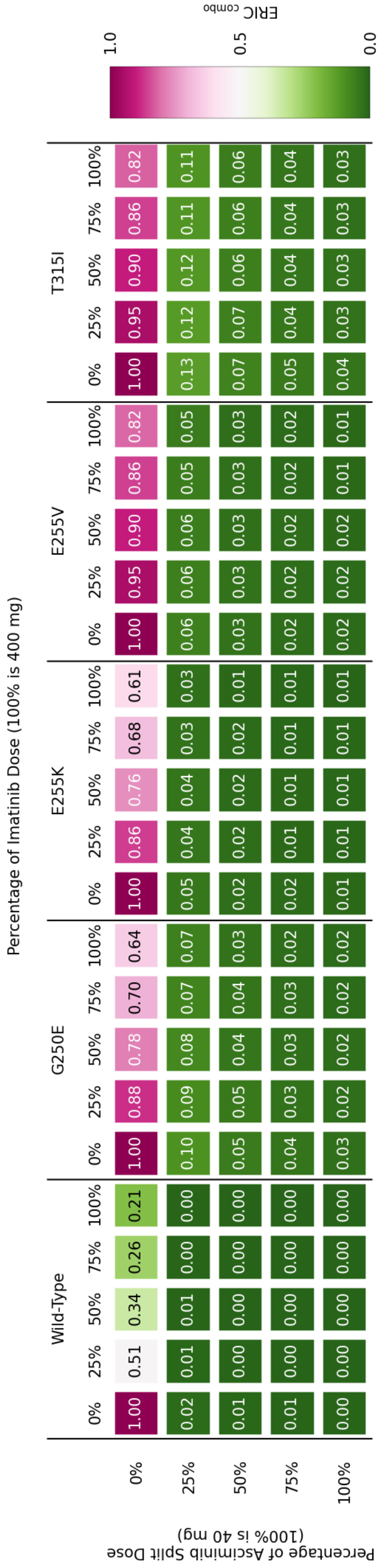


(b)

Figure S7: $ERIC_{combo}$ values for all combinations of mutations and doses for imatinib with (a) once-daily and (b) twice-daily asciminib.

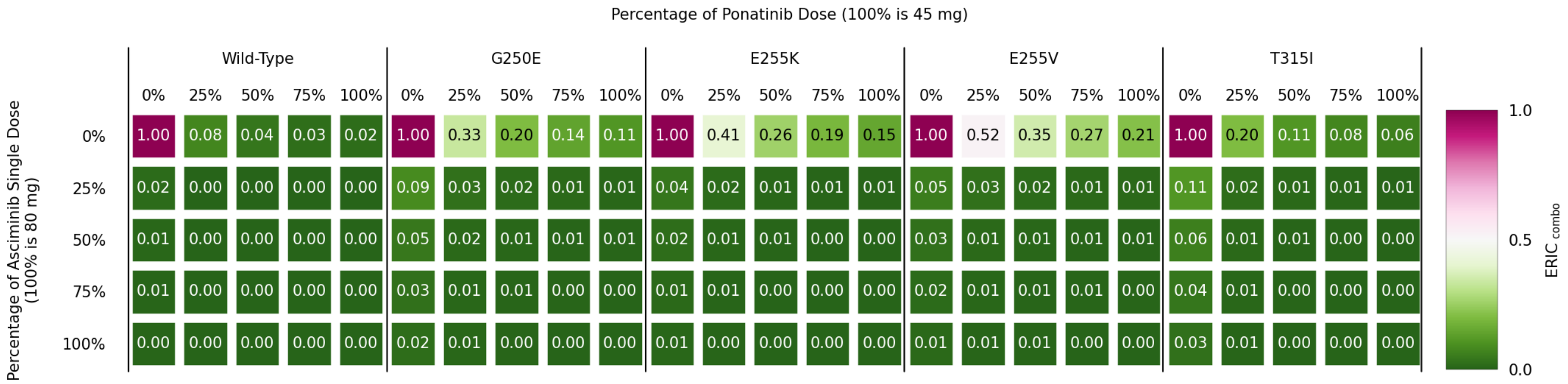


(a)

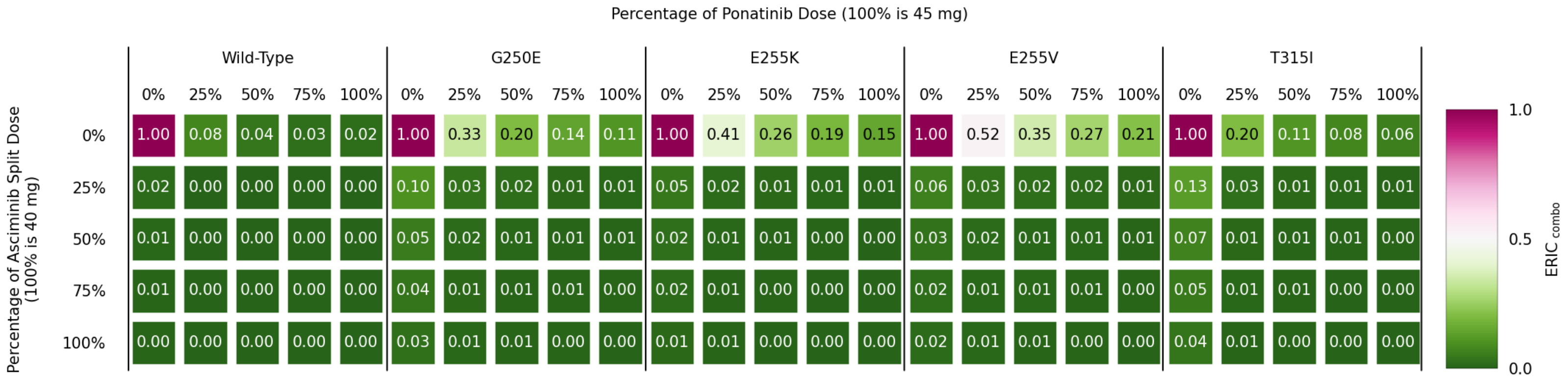


(b)

Figure S8: $ERIC_{combo}$ values for all combinations of mutations and doses for ponatinib with (a) once-daily and (b) twice-daily asciminib.

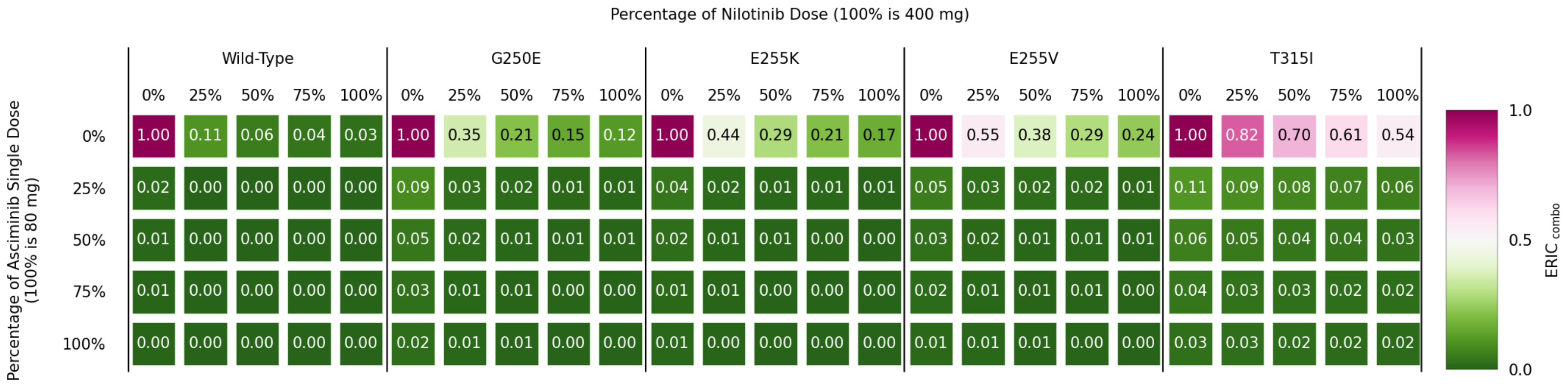


(a)

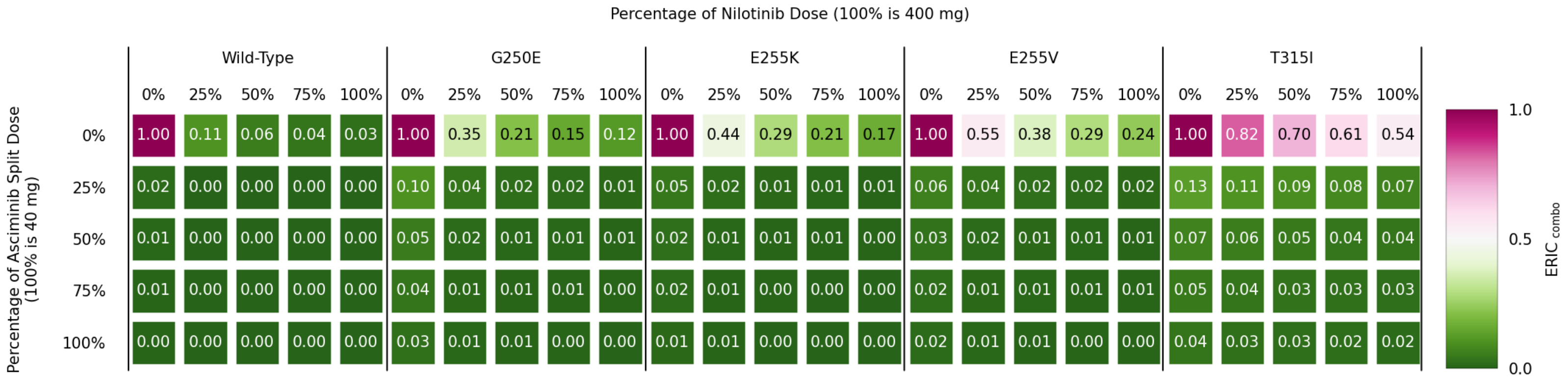


(b)

Figure S9: $ERIC_{combo}$ values for all combinations of mutations and doses for nilotinib with (a) once-daily and (b) twice-daily asciminib.

#### Examination of synergy

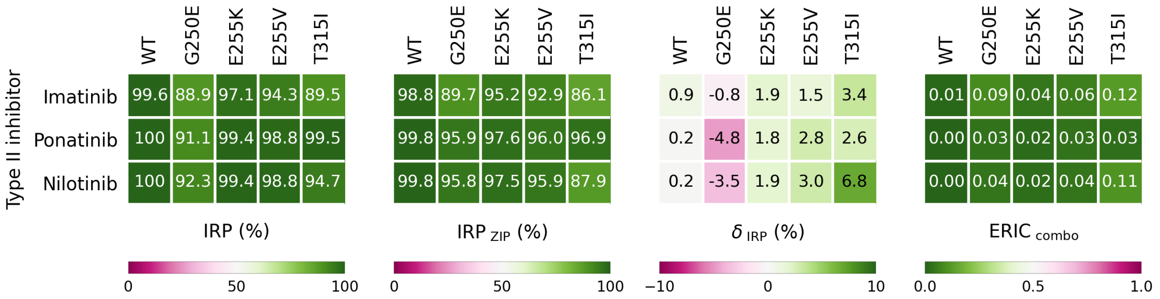


Figure S10: Comparison of IRP, $IRP_{ZIP}$, $\delta_{ZIP}$, and the product of Effective Ratios of $IC_{50}$ ($ERIC_{combo}$) for 25% dose of type II inhibitors and 25% dose of twice-daily asciminib. An equivalent figure for the once-daily asciminib regime can be found in Figure 8 in the main text.

### Full $\mathrm{IRP}_{\mathrm{ZIP}}$ and $\delta_{\mathrm{IRP}}$ heat maps

In the main text in Figure 8 and in supporting information Figure S10, we present an overview of the calculated $\mathrm{IRP}_{\mathrm{ZIP}}$ and $\delta_{\mathrm{IRP}}$ values, here we show the values calculated for every combination of dose, mutation, and inhibitor.

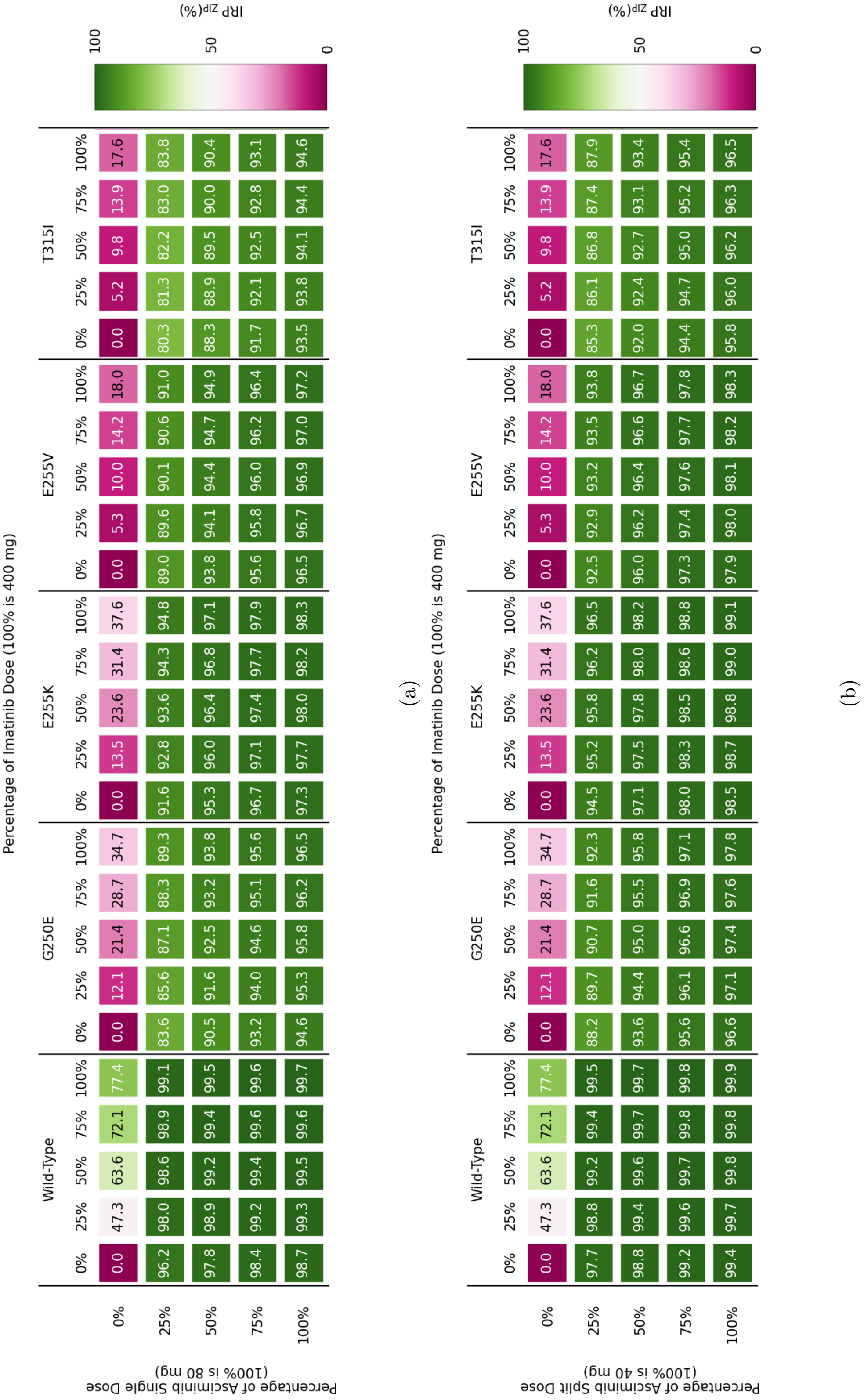


Figure S11: $\mathrm{IRP}_{\mathrm{ZIP}}$ values for all combinations of mutations and doses for imatinib with (a) once-daily and (b) twice-daily asciminib.

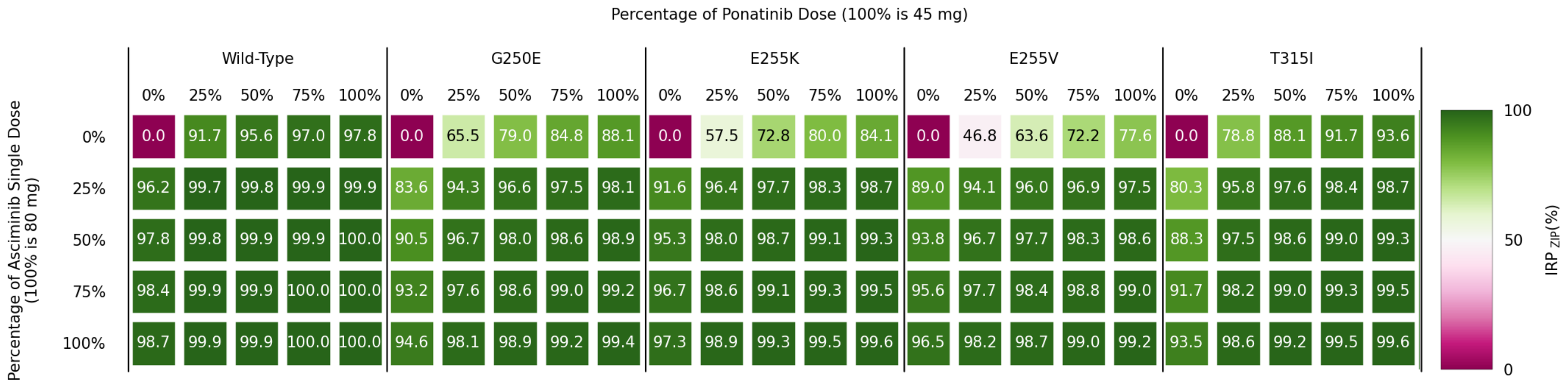


(a)

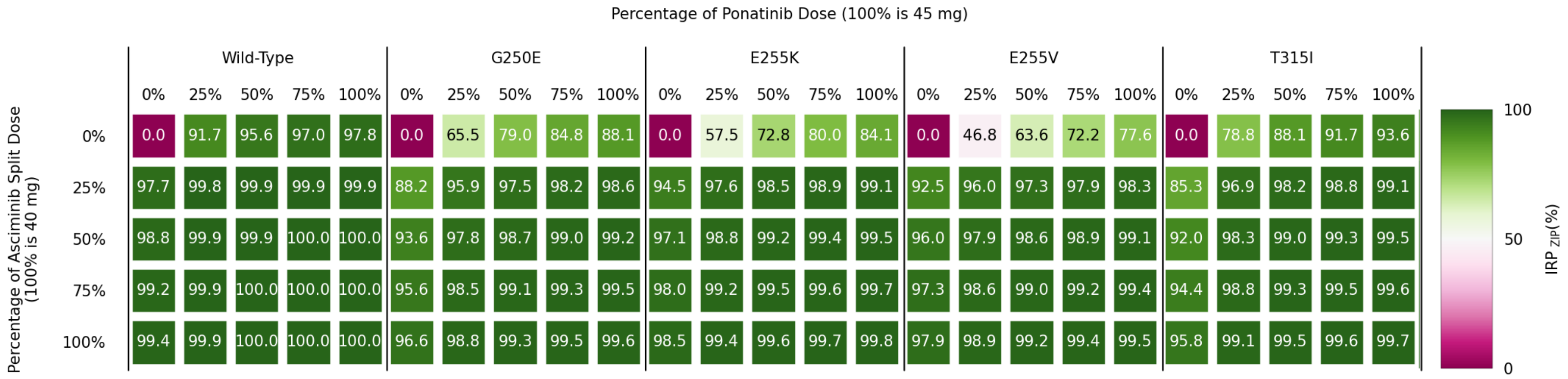


(b)

Figure S12: $IRP_{ZIP}$ values for all combinations of mutations and doses for ponatinib with (a) once-daily and (b) twice-daily asciminib.

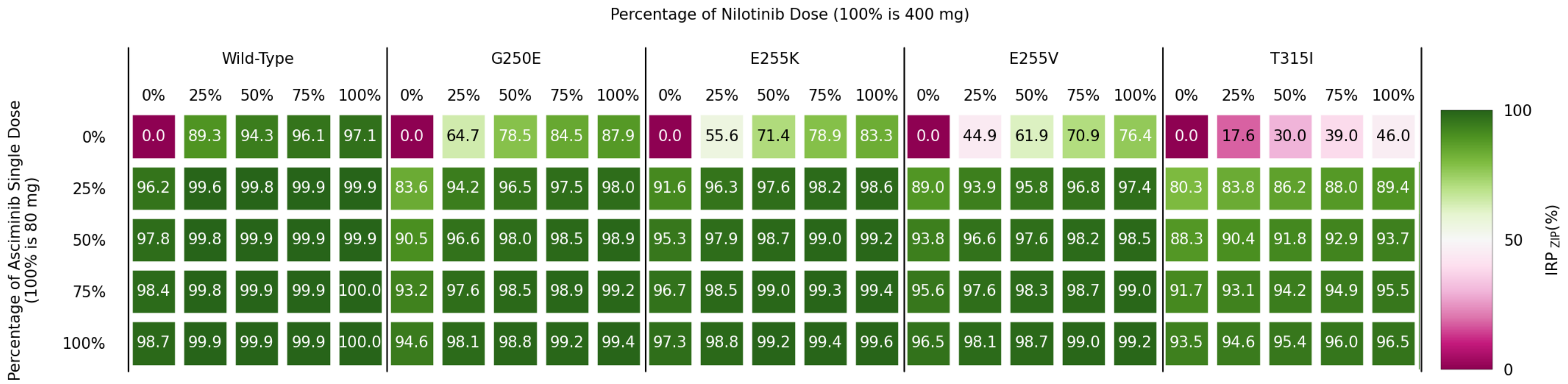


(a)

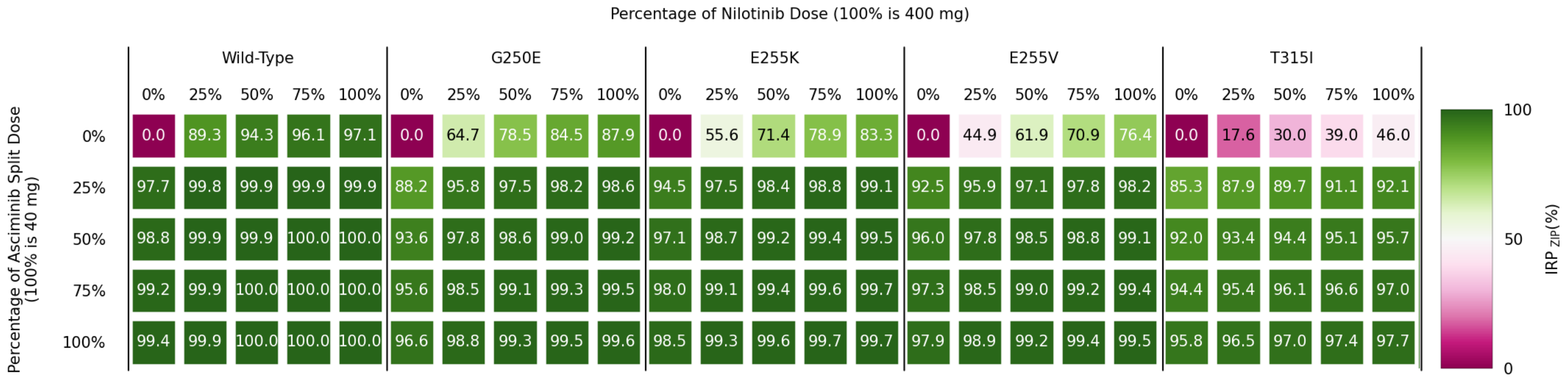


(b)

Figure S13: $IRP_{ZIP}$ values for all combinations of mutations and doses for nilotinib with (a) once-daily and (b) twice-daily asciminib.

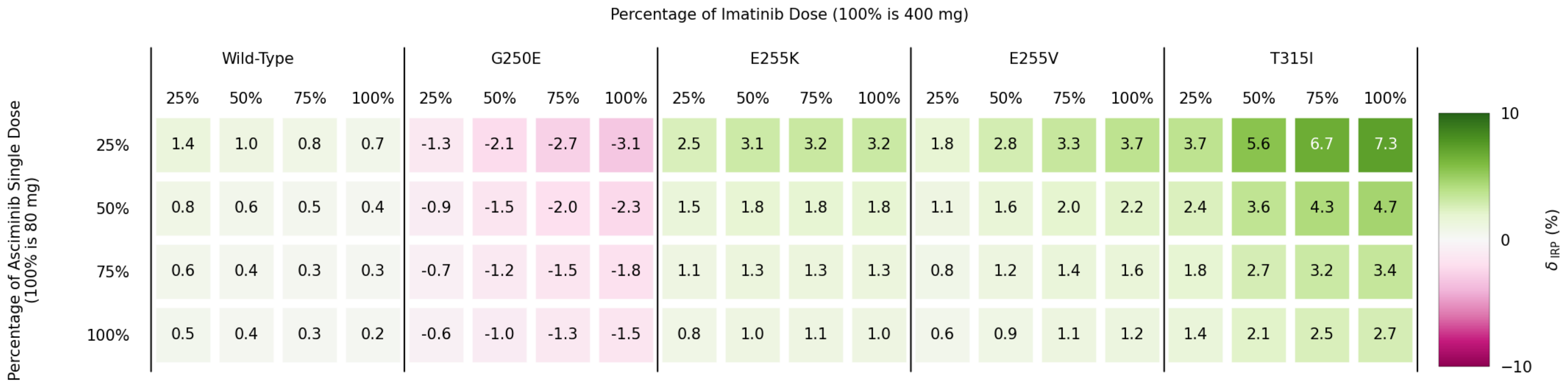


(a)

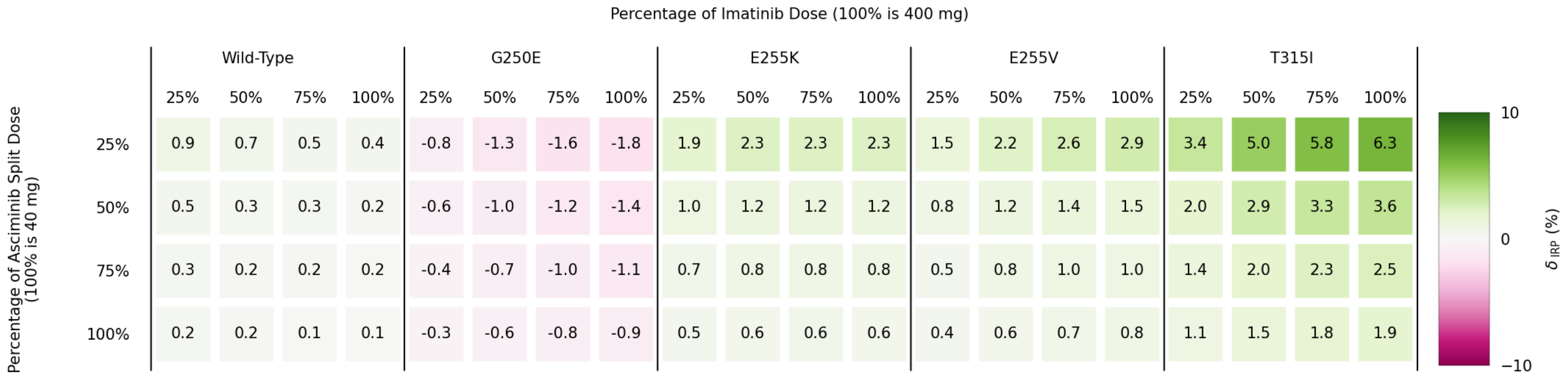


(b)

Figure S14: The $\delta_{\mathrm{IRP}}$ values for all combinations of mutations and doses for imatinib with (a) once-daily and (b) twice-daily asciminib.

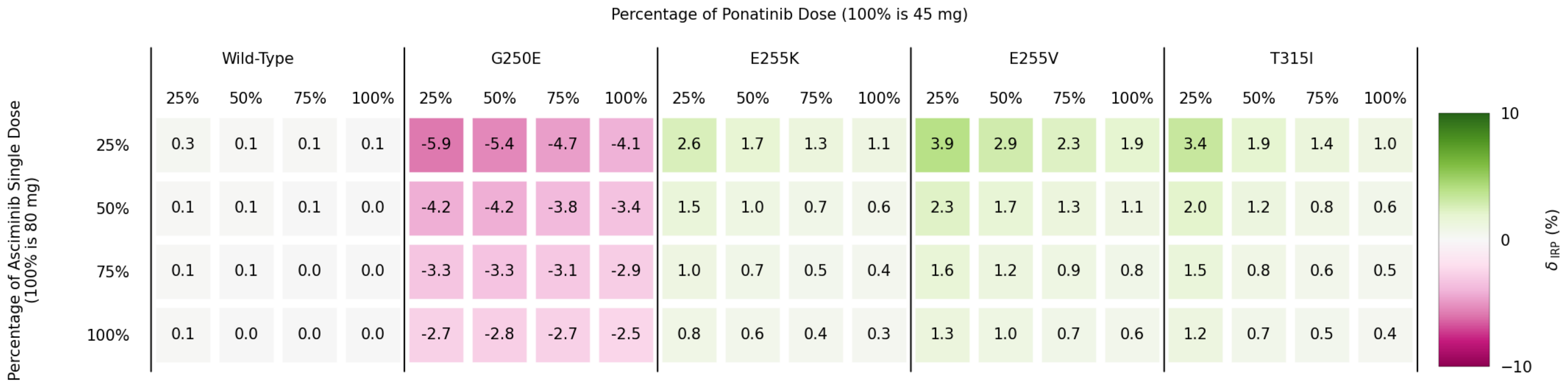


(a)

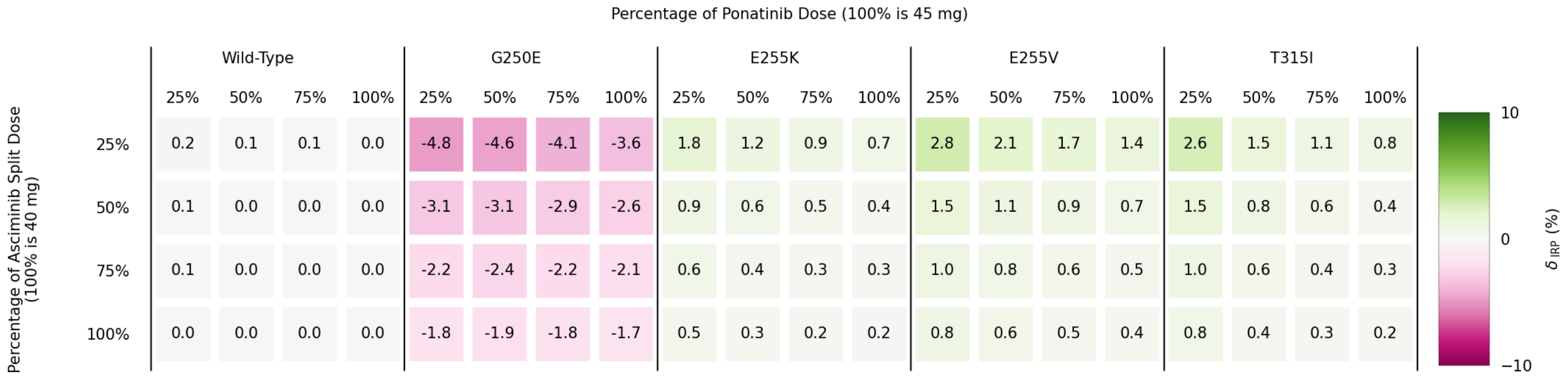


(b)

Figure S15: The $\delta_{\mathrm{IRP}}$ values for all combinations of mutations and doses for ponatinib with (a) once-daily and (b) twice-daily asciminib.

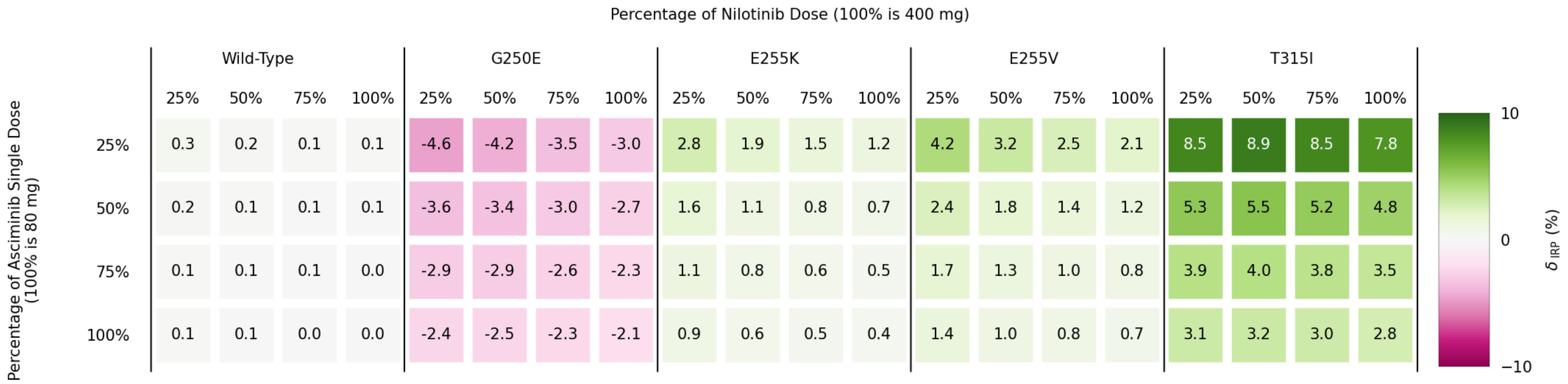


(a)

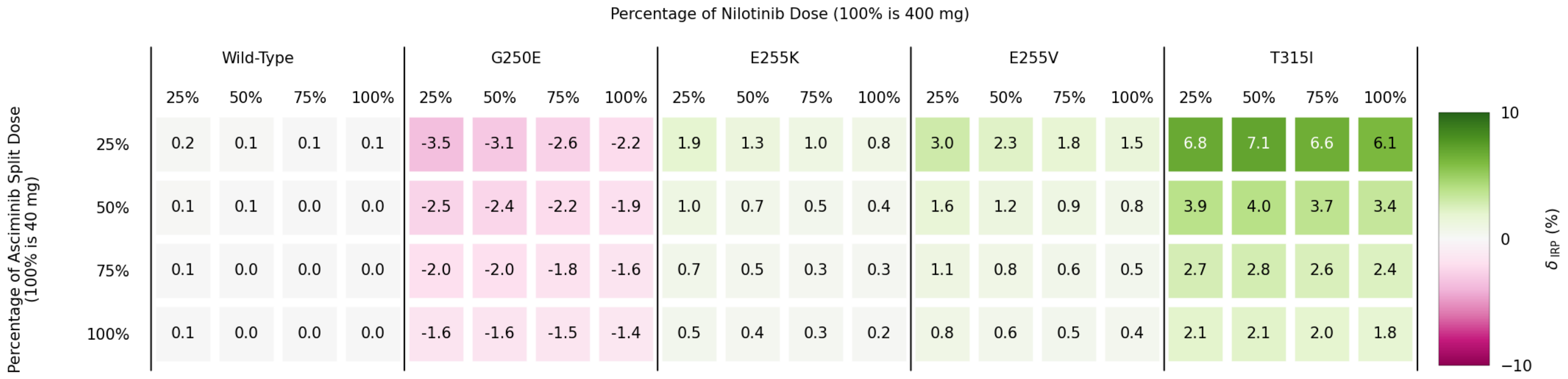


(b)

Figure S16: The $\delta_{\mathrm{IRP}}$ values for all combinations of mutations and doses for nilotinib with (a) once-daily and (b) twice-daily asciminib.

#### Selection of $k_{\mathbf{off}}^{\mathbf{Asc}}$

To select an appropriate $k_{\text{off}}^{\text{Asc}}$ value for the model, a range of values were inputted into the model of similar size to the unbinding rates of the type II inhibitors. We performed this for the WT and two mutants over a simulated 5 day period. The product rates did not vary much (see Figure S17), so $k_{\text{off}}^{\text{Asc}} = 0.01$ s was chosen.

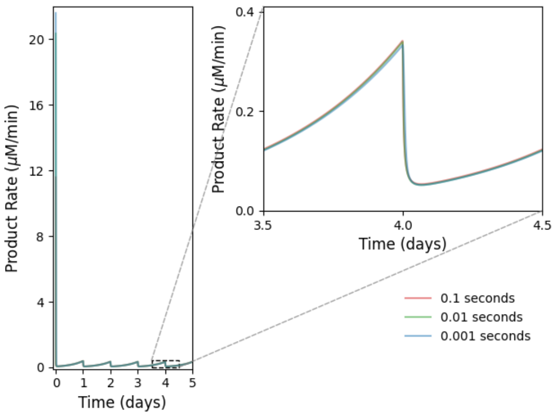


(a) Wild-type

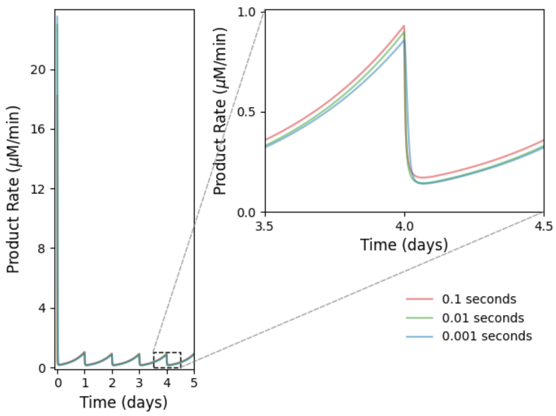


(b) E255K

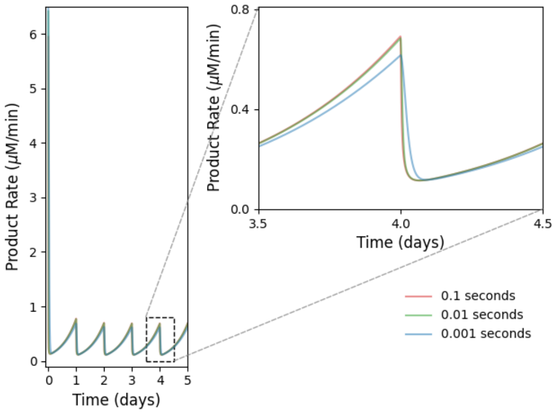


(c) T315I

Figure S17: Outputs of the product rate for models with different $k_{\text{off}}^{\text{Asc}}$ values.

#### Asciminib resistant mutants

Here we investigate three mutants that are associated to asciminib resistance: A337V, I502L and P465S. There are two reasons we do not deem it appropriate for these results to be directly compared to those outlined in the main text for the type II inhibitor resistant mutants.

1. The source of the $IC_{50}$ data for the mutants with associated asciminib resistance had very different $IC_{50}$ values for the WT for asciminib and the type II inhibitors than the data used in the main text.
2. We lacked other non-inhibitor related data for these mutants, so they have been assigned the corresponding values from the WT enzyme.

The data used in this version of the model can be seen in Tables S2 and S3, these values were derived in the same manner as those in the main text. Any values not appearing in these tables are equal to those for the WT in the main text. Examples of the state responses to different combinations can be found in Figure S18 and the IRP tables (similar to Figures S3 to S5) for this version of the model, can be found in Figures S19 to S21).

| Mutant | $IC_{50}$ (nM) | | | $K_R$ (nM) | | | $k_{on}^{R}$ $\times 10^{-4}(\text{min}^{-1})$ | | |
|---|---|---|---|---|---|---|---|---|---|
| | I | P | N | I | P | N | I | P | N |
| WT | 90.5 | 0.37 | 3.52 | 288.9 | 1.181 | 11.24 | 2.0423 | 41.32 | 17.80 |
| A337V | 82.1 | 3.52 | 3.67 | 262.1 | 11.24 | 11.72 | 2.251 | 4.343 | 17.07 |
| I502L | 56.2 | 0.29 | 2.68 | 179.4 | 0.9257 | 8.555 | 3.289 | 52.72 | 23.38 |
| P465S | 92.2 | 3.64 | 3.30 | 294.3 | 11.62 | 10.53 | 2.005 | 4.200 | 18.99 |

Table S2: Input values for imatinib (I), ponatinib (P), and nilotinib (N) for the asciminib resistant mutants. $IC_{50}$ values from source 1.

| Mutant | $IC_{50}$ (nM) | $K_A^I$ | $K_A^A$ | $k_I^{on}$ $\times 10^{-4}(\text{min}^{-1})$ | $k_A^{on}$ $\times 10^{-4}(\text{min}^{-1})$ |
|---|---|---|---|---|---|
| WT | 0.61 | 0.511 | 1.01 | 195.7 | 99.23 |
| A337V | 453 | 379 | 748 | 0.2635 | 0.1336 |
| I502L | 30.2 | 25.3 | 49.9 | 3.953 | 2.004 |
| P465S | 369 | 309 | 610 | 0.3235 | 0.1640 |

Table S3: Input values for asciminib resistant mutants. $IC_{50}$ values from source 1.

It appears that the co-binding is a major influence in the system response with these mutants as with those in the main text; however, resistance associated with asciminib appears to be to a lesser degree than the resistance associated with type II inhibitors.

Source 1 - Paul W. Manley, Louise Barys, Sandra W. Cowan-Jacob, The specificity of asciminib, a potential treatment for chronic myeloid leukemia, as a myristate-pocket binding ABL inhibitor and analysis of its interactions with mutant forms of BCR-ABL1 kinase, *Leukemia Research*, 98, 2020 (doi.org/10.1016/j.leukres.2020.106458)

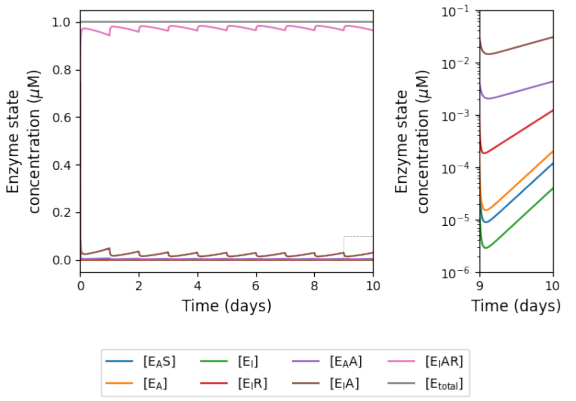


(a) WT with 25% imatinib and 25% once-daily asciminib.

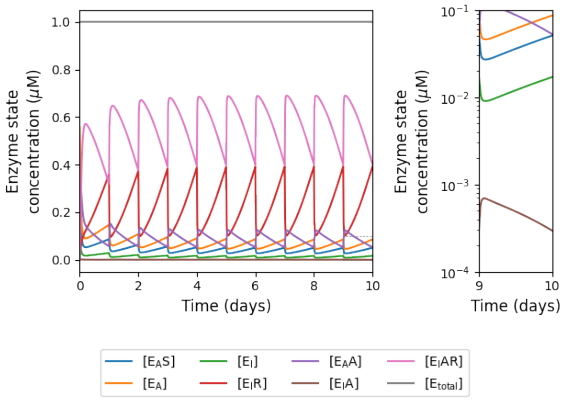


(b) A337V with 25% ponatinib and 25% once-daily asciminib.

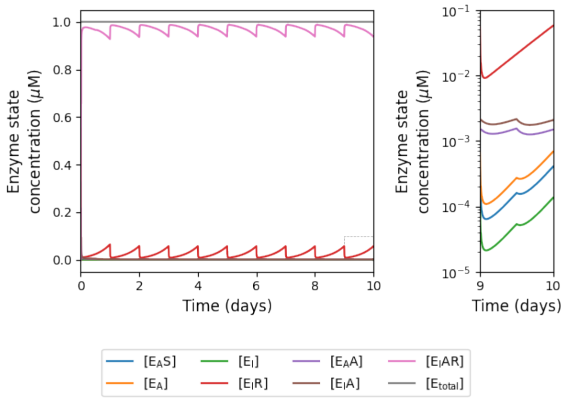


(a) I502L with 25% nilotinib and 25% once-daily asciminib.

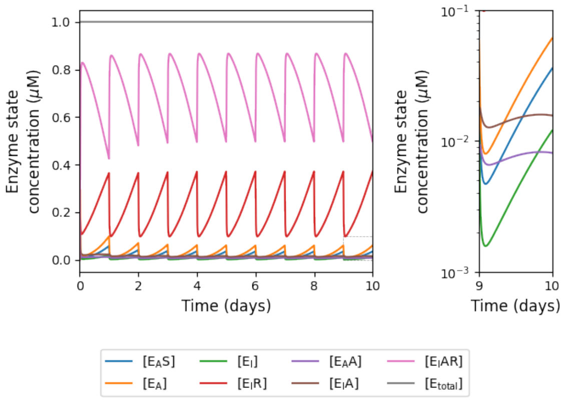


(b) P465S with 25% imatinib and 25% once-daily asciminib.

Figure S18: A sample of the effect on enzyme states under different doses for different mutations.

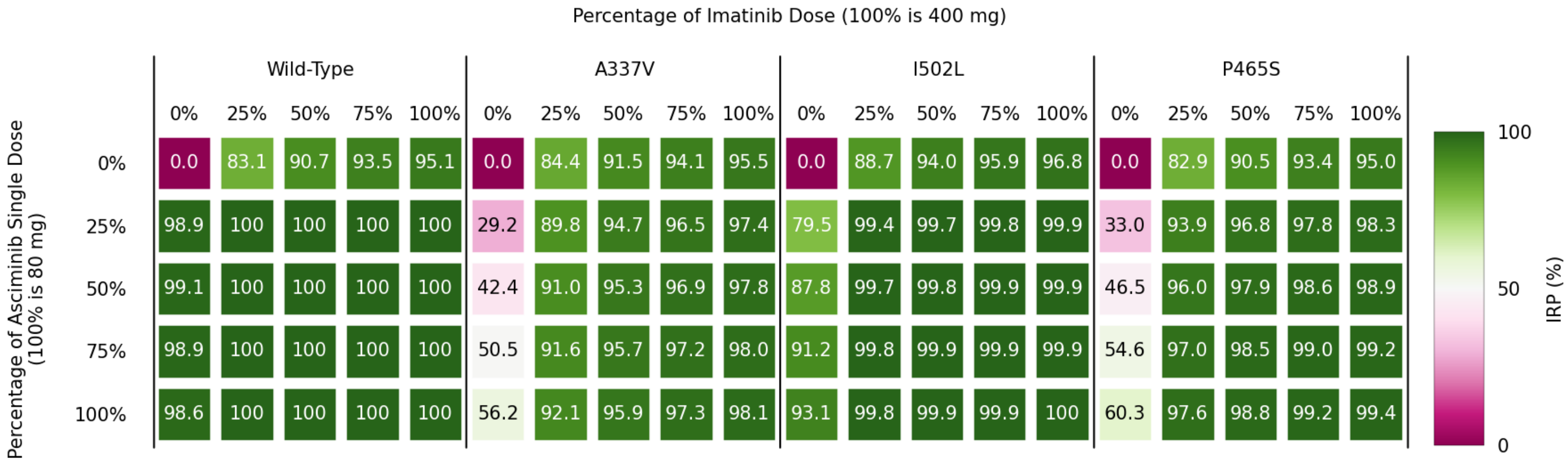


Figure S19: IRP of combinations of asciminib resistant mutations and doses for imatinib with once-daily asciminib regime.

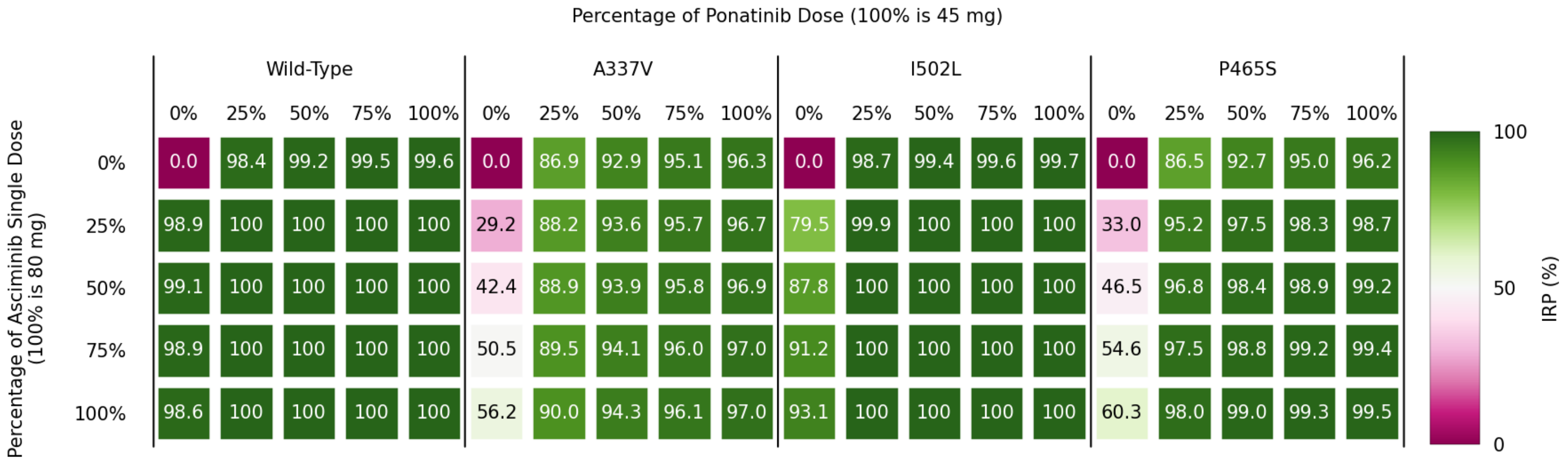


Figure S20: IRP of combinations of asciminib resistant mutations and doses for ponatinib with once-daily asciminib regime.

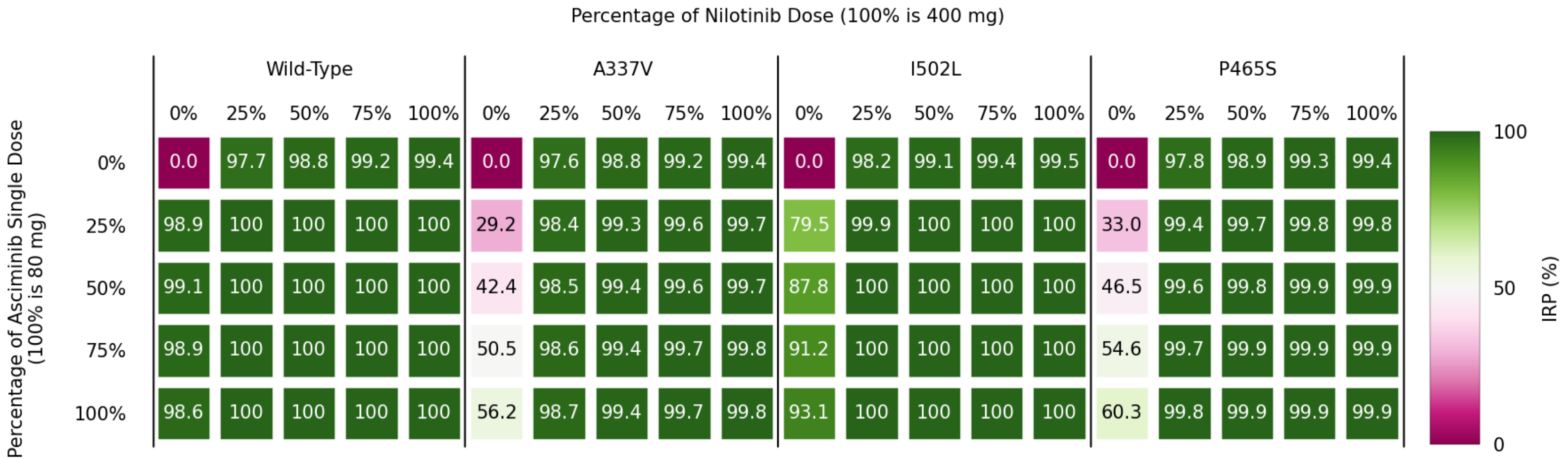


Figure S21: IRP of combinations of asciminib resistant mutations and doses for nilotinib with once-daily asciminib regime.
